# Increase of permissive secondary antiviral mutations in the evolution of A(H1N1)pdm09 influenza virus neuraminidases

**DOI:** 10.1101/2024.06.18.599480

**Authors:** Susanne Duwe, Jeanette Milde, Alla Heider, Marianne Wedde, Brunhilde Schweiger, Ralf Dürrwald

## Abstract

**Background:** The unexpected emergence of oseltamivir-resistant A(H1N1) viruses in 2008 was facilitated in part by the establishment of permissive secondary neuraminidase (NA) substitutions that compensated for the fitness loss due to the NA-H275Y resistance substitution. These viruses were replaced in 2009 by oseltamivir-susceptible A(H1N1)pdm09 influenza viruses.

**Methods:** Genetic analysis and screening A(H1N1)pdm09 viruses circulating in Germany between 2009 and 2024 were conducted to identify any potentially permissive or resistance-associated NA substitutions. Selected viruses were then subjected to further characterization in vitro.

**Results:** In the NA gene of circulating A(H1N1)pdm09 viruses, two secondary permissive substitutions, NA-V241I and NA-N369K, were identified. These substitutions demonstrated a stable lineage in phylogenetic analysis since the 2010-2011 influenza season. The data indicates a slight increase in viral NA bearing two additional potentially permissive substitutions, NA-I223V and NA-S247N, in the 2023-2024 season, that both result in a slight reduction in susceptibility to NA inhibitors.

**Conclusions:** The accumulation of secondary permissive substitutions in the NA of A(H1N1)pdm09 viruses increases the probability of the emergence of antiviral-resistant viruses. Therefore, it is crucial to closely monitor the evolution of circulating influenza viruses and to develop additional antiviral drugs against different target proteins.

## 1. Introduction

Influenza remains one of the most important infectious diseases in the world, causing seasonal epidemics with an estimated 3 to 5 million severe cases and 300,000 to 500,000 deaths worldwide each year [2]. In Germany, influenza A and B viruses typically affect 5-10% of adults and 20-30% of children, causing millions of acute respiratory infections.At the time of the 2019 coronavirus (COVID-19) pandemic, influenza viruses were almost not circulating globally [1, 3]. Circulation of influenza viruses returned very early in comparison of pre-pandemic influenza seasons, after the non-pharmaceutical interventions (NPIs) such as masks and improved hand hygiene were eased. In Germany, the 2022-2023 influenza season peaked in December 2022 and was dominated by A(H3N2) viruses, while the 2023-2024 season, dominated by A(H1N)pdm09 viruses peaked in comparison to pre-pandemic seasons a few weeks earlier (in preparation).

Although NPIs can be effective in preventing influenza infection, particularly in clinical or household settings, vaccination is considered the most effective method of prevention [4]. Clinical efficacy is influenced by the circulating viruses, the vaccination regimen, the adjuvant used, the duration of immunity induced by the particular vaccine virus, and the immune status of the vaccinee, which is determined by age, previous antigenic exposure to influenza viruses, immunologic reactivity, and the timing of vaccination. Antiviral prophylaxis and treatment are additional options in situations where vaccination is not feasible, such as in vulnerable patient populations with reduced vaccine efficacy or the emergence of a new virus subtype [5, 6]. Its use should be considered in patients at high risk for severe complications of influenza and in critically ill hospitalized patients, according to most international and national public health organizations and medical and scientific societies [7].

The two neuraminidase inhibitors oseltamivir and zanamivir and the polymerase inhibitor baloxavir marboxil are currently recommended in Europe for the prevention and treatment of influenza. The adamantanes rimantadine and amantadine should no longer be used for the prophylaxis and treatment of influenza infections due to the natural polymorphism in the M2 ion channel, which confers resistance of influenza viruses [7].

Since baloxavir is only available through international pharmacies in Germany, zanamivir must be administered either by inhalation or intravenously, adamantane is not effective against influenza B, and circulating influenza A viruses are resistant, the orally available oseltamivir remains the drug of choice 25 years after its first approval by the European Medicines Agency (EMA) [7, 8].

Resistance to neuraminidase inhibitors has occurred only sporadically since their approval and mainly in clinical trials in immunocompromised patients. Reduced susceptibility to oseltamivir, caused by the exchange of histidine for tyrosine at position 275 (H275Y) of the neuraminidase of seasonal A(H1N1) viruses, has been particularly common. In the absence of drugs, the mutations that confer antiviral resistance may have a detrimental effect on viral fitness, so secondary fitness-restoring mutations must occur to allow resistance to spread on a large scale [9]. Until November 2007, the prevalence of resistant A(H1N1) viruses in untreated adults was <1%. The increased circulation of oseltamivir-resistant A(H1N1) viruses in treatment-naive patients in the northern hemisphere observed in winter 2007-2008 was unexpected. These viruses remained susceptible to zanamivir. The prevalence of oseltamivir-resistant viruses increased sharply during the season, with an average of one in four A(H1N1) viruses resistant to oseltamivir due to NA-H257Y substitution, circulating in Europe during the 2007-2008 season. In the following seasons, 2008 in the southern hemisphere and 2008-2009 in the northern hemisphere, all A(H1N1) viruses analyzed showed an unchanged resistance profile and remained stable in their properties. Phylogenetic analysis revealed two amino acid substitutions in the neuraminidase protein that appeared to anticipate NA-H275Y - the NA-V234M and NA-R222Q. Several fitness experiments confirmed that these NA substitutions restore the fitness of H275Y-bearing viruses compared to that observed in wild-type viruses [10]. NA-H275Y decreases the amount of neuraminidase that reaches the cell surface which is of disadvantage for viral replication; the permissive secondary antiviral mutations can counteract this process and restore full viral replication.

These mutations were defined as permissive secondary antiviral mutations that do not confer resistance but support fitness of influenza viruses bearing the antiviral mutation NA-H275Y. These substitutions occurred in the influenza viruses just prior to the widespread emergence of oseltamivir-resistant A(H1N1) viruses carrying NA-H275Y. Although the permissive secondary substitutions NA-V234M and NA-R222Q are located distantly from NA-H275Y they are prerequisites for the emergence of fit resistant H275Y viruses because they maintain adequate surface NA expression which otherwise would be disrupted by NA-H275Y [9, 10].

With the beginning of the influenza pandemic in June 2009, seasonal A(H1N1) viruses were replaced by viruses of the A(H1N1)pdm09 subtype. Due to reassortment events, these A(H1N1)pdm09 viruses carry a neuraminidase from swine influenza A viruses that is characterized by susceptibility to neuraminidase inhibitors [11-14]. Since then frequency of circulating resistant A(H1N1)pdm09 remains low (<1%) [15], but computational and community outbreak analyses have provided evidence for potential permissive secondary NA substitutions that alter the viral fitness of oseltamivir-resistant A(H1N1)pdm09 virus strains [5]. In this context, a compensatory role by conferring a robust fitness on viruses bearing substitutions NA-V241I and NA-N369K, which correlated with enhanced surface expression and enzymatic activity of the A(H1N1)pdm09 NA protein has been demonstrated [16-18].

In addition, the NA-I223V/R and NA-S247N substitutions have been shown to potentially increase the resistance of A(H1N1)pdm09 viruses to neuraminidase inhibitors in reverse genetics and in vitro analyses [5, 19].

The aim of our study was to monitor the evolution of these four secondary permissive substitutions, to assess their impact on the viral fitness of A(H1N1)pdm09 viruses in the context of the continuous monitoring of antiviral resistance of influenza viruses circulating in Germany.

## 2. Materials and Methods

### Clinical specimens and influenza virus typing and subtyping

Medical practices participating in the national sentinel surveillance system collected clinical specimens from outpatients presenting with acute respiratory or influenza-like illness (nasal, throat or pharyngeal swabs) and sent them in viral transport media to the German National Influenza Centre (NIC) at the Robert Koch Institute. The Ethics Committee of the Charité University Hospital in Berlin (reference EA2/126/11) gave written permission for the German national surveillance of influenza and other respiratory viruses. Influenza sentinel surveillance is regulated by German legislation (§13, §14, Infektionsschutzgesetz). All analyses were conducted pseudonymised. All sentinel patients gave written informed consent.

The swabs collected between seasons 2009/2010 and 2023/2024 were washed in cell culture medium (MEM/HEPES, Minimum Essential Media with Hepes and 1% penicillin/streptomycin) and viral RNA was extracted. Complementary DNA was synthesized by random reverse transcription (Invitrogen, Carlsbad, California). For in-house multiplex typing and subtyping of influenza virus, primers and probes targeting the M, HA, and NA genes were used, as previously described [4, 20].

### Viral propagation

Influenza viruses were propagated in cell culture using MDCK-SIAT 1 cells (European Collection of Cell Cultures (ECACC, Salisbury, United Kingdom), Lot: 05G023) by inoculating cell monolayers with sterile filtered swab suspension as described recently [5, 21].

### Neuraminidase inhibition test

Oseltamivir carboxylate and zanamivir was kindly provided free of charge by Hoffmann-La Roche Ltd. (Basel, Switzerland) and by GSK plc. (London, United Kingdom), respectively. Stock solutions (100 µM in sterile ultrapure double-distilled water) were stored at -20°C and for susceptibility testing diluted in MES buffer [32.5 mM morpholine ethanesulfonic acid (Sigma-Aldrich), pH 6.5, and 4 mM CaCl2]. Neuraminidase inhibition test was measured using 2’-(4-methylumbelliferyl)-α-d-N-acetylneuraminic acid (Munana; Biosynth AG, Staad SG, Switzerland and Sigma-Aldrich, St. Louis, Missouri, USA) as substrate, as described previously [22]. The emitted fluorescence values of the released 4-methylumbelliferone were measured in a spectrofluorometer (Tecan AG, Männedorf, Switzerland). The 50% inhibitory concentration (IC_50_) was calculated as mean of the 50% inhibitory concentration ± standard deviation (IC_50_ ± SD) of duplicate to quadruplicate assays from the dose-response curve using MS Excel software (MS Office 2010) and compared to the reference IC_50_ values [12]. Neuraminidase susceptibility were judged following WHO recommendations, that defined reduced and highly reduced susceptibility to NAIs by a ⩾ 10- to 100-fold and >100-fold increase in the NAI IC_50_ compared to the NAI IC_50_ of the sensitive control (a wild-type virus of the same type or subtype) [23].

### Enzyme kinetic

Neuraminidase activity was determined by using MUNANA fluorogenic substrate as described above. The final concentration of the substrate ranged from 0.72 µM to 509 µM. Michaelis–Menten constants (Km) were calculated by using the Lineweaver–Burk diagrams generated with Excel software (Microsoft) [22].

### Genome sequencing and resistance analysis

Influenza virus RNA was extracted, viral genome amplified by specific PCR. PCR products of amplified NA segments were sequenced by automated nucleotide cycle sequencing (primer sequence on request) using the BigDye®Terminator v3.1Cycle Sequencing Kit (Applied Biosystems, Darmstadt, Germany) and a capillary sequencer 3130xl (Applied Biosystems) or underwent whole/next generation sequencing as described previously [5, 24, 25].

The GISAID accession numbers of sequences analyzed are listed in Supplementary Table 1, which is part of the supplementary data provided with this manuscript. NGS data were analyzed for molecular resistance markers in neuraminidase by FluSurver enabled by data from GISAID (https://flusurver.bii.a-star.edu.sg/).

**Table 1.** Prevalence of permissive substitutions and susceptibility to NA inhibitors of circulating viruses calculated as means of 50% inhibitory concentration (IC50) between April 2009 and April 2024.

| Influenza season | Prevalence of substitutions % (detected/tested) |  |  |  |  | X IC<sub>50</sub> ± SD (nM) |  |  |
| --- | --- | --- | --- | --- | --- | --- | --- | --- |
|  | NA-223V | NA-247N | NA-241I | NA-369K | N<sup>1</sup> | oseltamivir | zanamivir | N<sup>2</sup> |
| 2009 (April-September) | 0 | 0 | 0 | 0 | 64 | 1.66 ± 0.52 | 0.39 ± 0.20 | 65 |
| 2009-2010 | 0 | 0 | 0 | 0 | 94 | 1.41 ± 0.63 | 0.38 ± 0.17 | 142 |
| 2010-2011 | 0 | 1.5<br>(1/65) | 66<br>(43/65) | 62<br>(40/65) | 65 | 1.19 ± 0.61 | 0.52 ± 0.83 | 65 |
| 2011-2012 | 0 | 0 | 100 | 100 | 11 | 2.15 ± 0.49 | 0.45 ± 0.07 | 2 |
| 2012-2013 | 0 | 0 | 100 | 100 | 9 | 1.63 ± 0.46 | 0.25 ± 0.09 | 31 |
| 2013-2014 | 0 | 0 | 100 | 100 | 17 | 1.85 ± 0.41 | 0.51 ± 0.33 | 15 |
| 2014-2015 | 0 | 0 | 100 | 100 | 25 | 0.94 ± 0.37 | 0.50 ± 0.18 | 147 |
| 2015-2016 | 0.05 (1/212)<br>223R | 0 | 100 | 100 | 212 | 0.90 ± 0.36 | 0.43 ± 0.33 | 147 |
| 2016-2017 | 0 | 0 | 100 | 100 | 4 | 1.68 ± 0.35 | 0.77 ± 0.18 | 9 |
| 2017-2018 | 0 | 0 | 100 | 100 | 57 | 1.13 ± 0.40 | 0.62 ± 0.24 | 147 |
| 2018-2019 | 0 | 0 | 100 | 100 | 103 | 0.97 ± 0.36 | 0.48 ± 0.30 | 144 |
| 2019-2020 | 0 | 0 | 100 | 100 | 162 | 1.02 ± 0.42 | 0.34 ± 0.14 | 145 |
| 2020-2021 | no viruses detected |  |  |  |  |  |  |  |
| 2021-2022 | 0 | 0 | 100 | 100 | 17 | 0.96 ± 0.35 | 0.30 ± 0.06 | 7 |
| 2022-2023 | 0 | 1.0<br>(1/86) | 100 | 100 | 86 | 0.82 ± 0.33 | 0.26 ± 0.10 | 85 |
| 2023-2024 | 0.7<br>(4/552) | 1.1 (6/552) | 100 | 100 | 552 | 0.90 ± 0.26 | 0.24 ± 0.10 | 145 |

X = mean of seasonal treshold, SD = standard deviation, nM = nanoMolar, N<sup>1</sup>= Number of sequences analyzed, N<sup>2</sup>= Number of isolates tested

### Phylogenetic analysis

Phylogenetic evaluation was done using Mega 7, neighbour-joining method, phylogeny test by bootstrap method and 1000 replications, Kimura 2 substitution model and partial deletion (site coverage cutoff 5 %) [24].

## 3. Results

### 3.1 Permissive secondary substitutions of (H1N1)pdm09virus neuraminidase NA-V241I, NA-N369K

Sequencing and phylogenetic analyses of 1478 A(H1N1)pdm09 NA genes collected in Germany between April 2009 and March 2024 indicate that the substitutions NA-V241I NA-369K first emerged during the 2010-2011 influenza season. Since then, viral NAs have been characterized by the V241I and N369K substitutions and form a cluster within the phylogenetic tree. (Figure 1, Table 1).

**Figure 1.**
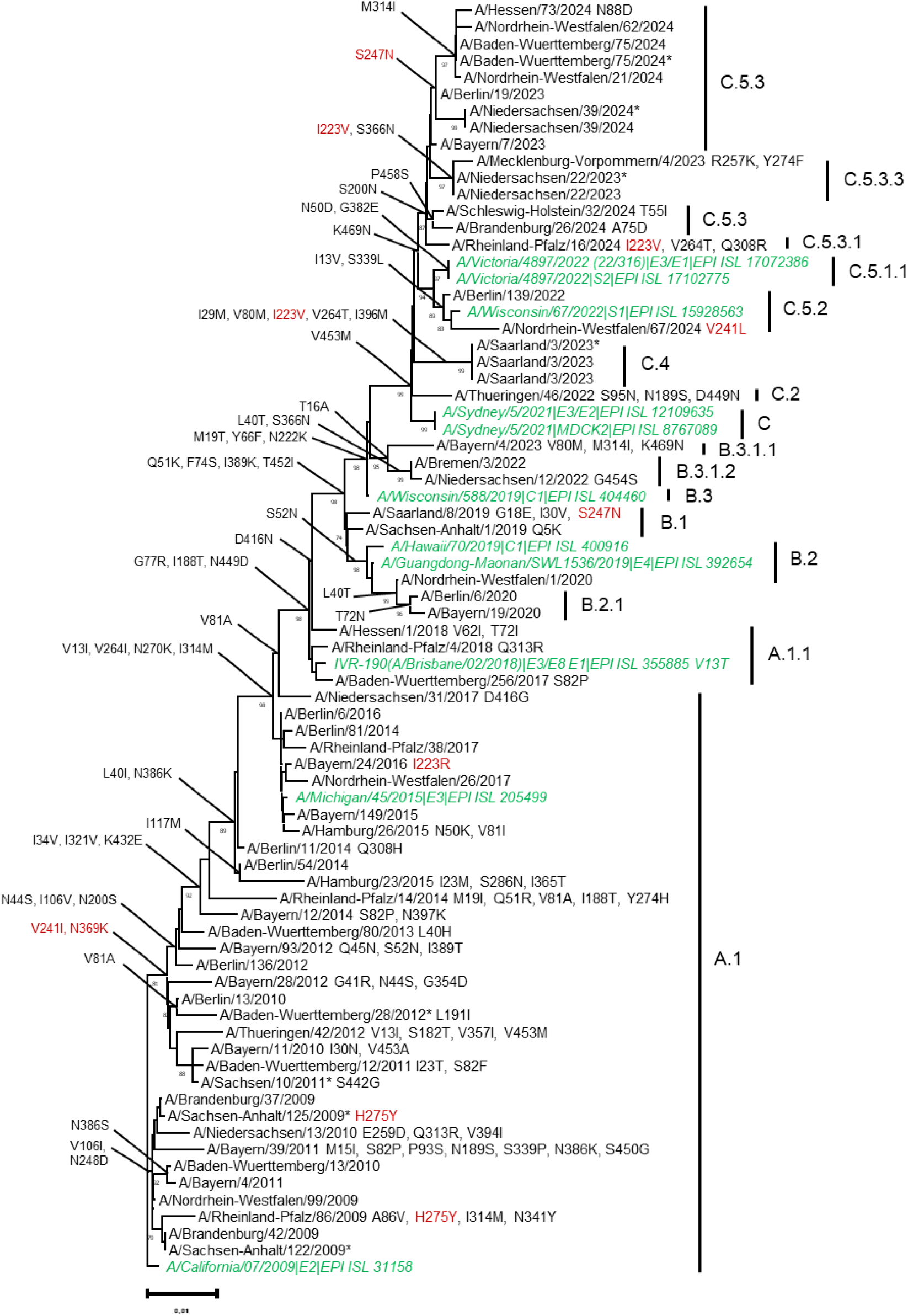
Phylogenetic analysis of A(H1N1)pdm09 neuraminidase genes from viruses circulating between 2009 and 2024 in Germany. sssssPhylogenetic analysis was performed by using cDNA sequences from A(H1N1)pdm09 neuramini-dase genes (nt 91-nt 1341). The amino acid substitution characterizing the respective NA-clade, that was calculated with https://clades.nextstrain.org/, are indicated. Phylogenetic evaluation was done using Mega 7, neighbor-joining method, phylogeny test by bootstrap method and 1000 replications, Kimura 2 substitution model and partial deletion (site coverage cutoff 5%). Reference strains from vaccine viruses were indicated in green, cell-cultured virus isolates are marked with an asterisk.

Comprehensive NA inhibitor susceptibility analyses of 1067 A(H1N1)pdm09 virus isolates collected between April 2009 and March 2024 in Germany using a fluorescence-based NA inhibition assay showed no correlation between the emergence of these potentially permissive NA mutations and NA inhibitor susceptibility (Table 1). The mean 50% inhibitory concentration expresses the susceptibility of viruses to the neuraminidase inhibitors oseltamivir and zanamivir and varies from season to season due to technical conditions. The NRC for influenza viruses complies with WHO recommendations and at least 20% of the influenza viruses detected in each season are tested for resistance. The number of viruses tested therefore reflects the circulation of A(H1N1)pdm09 viruses in each season. With a few exceptions, mainly due to previous treatment with oseltamivir, almost all viruses were susceptible to the NA inhibitors oseltamivir and zanamivir, giving a prevalence of oseltamivir-resistant viruses <1% (data not shown). The detected resistance to oseltamivir was due to the NA-H275Y substitution, which is known to cause strong resistance to oseltamivir; the viruses remain susceptible to zanamivir as shown using a reference virus in Table 2.

**Table 2:**
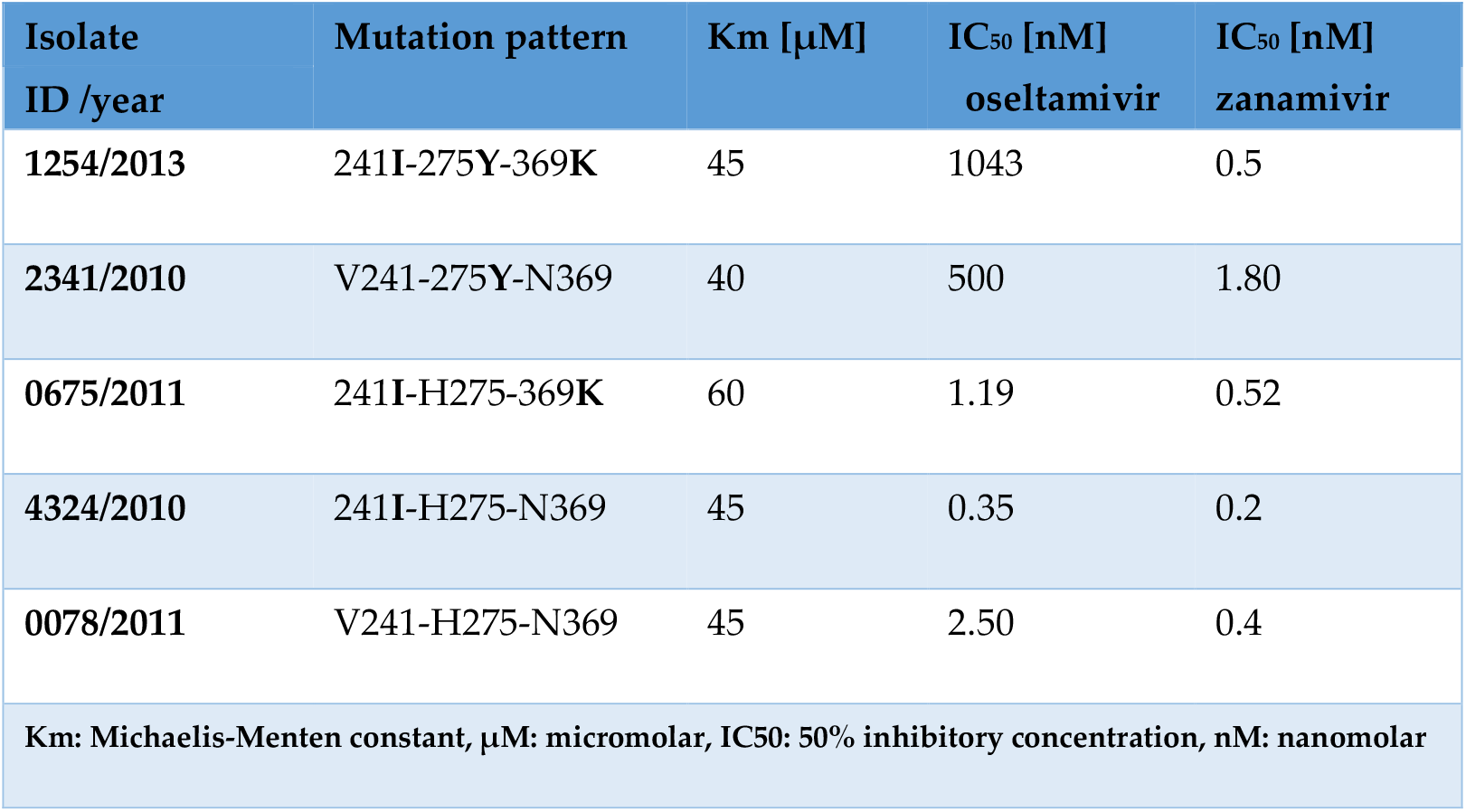
Enzymatic properties and antiviral susceptibility of A(H1N1)pdm09 NA.

| Isolate ID /year | Mutation pattern | Km [μM] | IC<sub>50</sub> [nM]<br>oseltamivir | IC<sub>50</sub> [nM]<br>zanamivir |
| --- | --- | --- | --- | --- |
| 1254/2013 | 241I-275Y-369K | 45 | 1043 | 0.5 |
| 2341/2010 | V241-275Y-N369 | 40 | 500 | 1.80 |
| 0675/2011 | 241I-H275-369K | 60 | 1.19 | 0.52 |
| 4324/2010 | 241I-H275-N369 | 45 | 0.35 | 0.2 |
| 0078/2011 | V241-H275-N369 | 45 | 2.50 | 0.4 |
| Km: Michaelis-Menten constant, μM: micromolar, IC<sub>50</sub>: 50% inhibitory concentration, nM: nanomolar | | | | |

To analyze the enzymatic properties of the A(H1N1)pdm09 virus NA, enzymes with different substitution patterns at positions 241, 275 and 369 were studied (Table 2). Determination of the Michaelis-Menten constant (Km) by NA enzyme kinetic assays shows a marginal increase in Km and therefore a slightly reduced enzyme affinity for its substrate in NA bearing NA-241I and NA-369K compared to wild-type NA-V241 and NA-N369. Together with the resistant substitution NA-H275Y, these two permissive NA substitutions showed no effect on enzyme affinity of oseltamivir resistant NA, suggesting a similar fitness of the virus strains with different mutation patterns (Table 2). On the other hand, the NA with the substitutions NA-241I and NA-369K together with NA-275Y had a 417-fold decreased susceptibility to oseltamivir compared to the wild-type *V*241-H275-N369. However, the susceptibility to oseltamivir of the NA-V241-275Y-N369 virus is only 200-fold reduced, suggesting a permissive effect of NA-241I and NA-369K on NA-275Y-mediated oseltamivir resistance.

### 3.2 Permissive secondary substitutions of (H1N1)pdm09virus neuraminidase NA-I223V, NA-S247N

In previous seasons, substitutions of the A(H1N1)pdm09 NA occurred only sporadically, but in the last seasons after the Covid-19 pandemic between 2022 and 2024, an increase of NA-223 and NA-247 substitutions from 0% in 2021-2022 to 1% in the 2023-2024 influenza season was observed using NGS-generated genome sequences. For further analysis, two NA-223V and four NA-247N A(H1N1)pdm09 viruses were grown in cell culture. The NA substitutions were confirmed in these cell-cultured viruses, indicating that they do not adversely affect viral fitness and are stable across multiple passages and virus generations. Analysis of the enzymatic properties of NA-223V showed in comparison to NA-I223 an only marginally decreased susceptibility to NA inhibitors demonstrated by IC_50_-analysis. The activity analysis of oseltamivir and zanamivir by IC_50_ determination showed decrease for NA-247N indicated by an up to 5-fold increase of inhibitor’s IC^50^ in comparison to that of wild-type NA-S247 (Table 3).

**Table 3:**
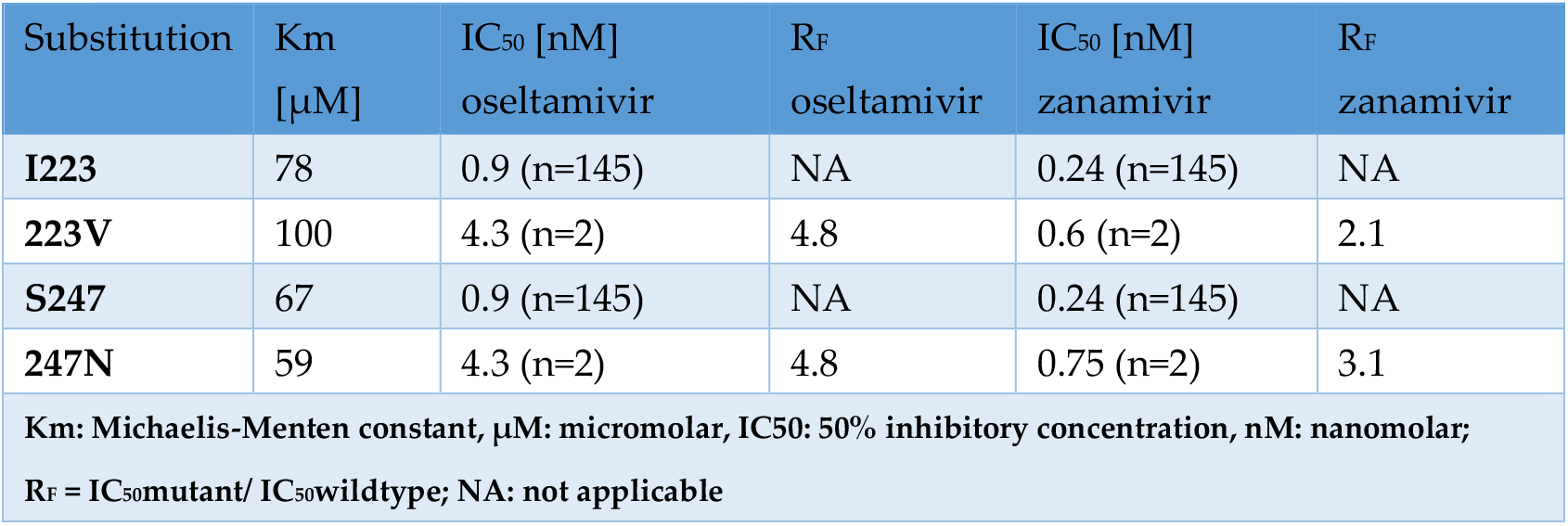
Enzyme affinity and susceptibility to NA-inhibitors of A(H1N1)pdm09-NA with and without permissive mutations calculated as Km and IC_50_ determination.

The enzymatic affinity of NA-223V to Munana substrate was characterized by a slight decrease of substrate affinity shown by an 1.3-fold increase of Km compared with wild-type NA-I223 (Table 3). The NA-S247N substitution does not affect the affinity of the enzyme, the Km of NA-247N was comparable to that of the wild-type NA-S247 (table 3).

## 4. Discussion

Neuraminidase inhibitors, primarily oral oseltamivir, remain the treatment of choice for influenza, especially in people who cannot be vaccinated and who are at risk of severe disease progression due to other medical conditions [7].

The emergence and rapid spread of oseltamivir-resistant NA-275Y A(H1N1) viruses in 2007 demonstrated that secondary supporting mutations can help to compensate for the loss of fitness caused by this resistance mutation. The NA-H275Y substitution in subtype N1 neuraminidase confers strong resistance to oseltamivir by reducing the amount of neuraminidase molecules that reach the cell surface. Phylogenetic analyses were used to identify substitutions that compensate for this loss of fitness [9, 10]. The NA-H222Q and NA-V234M substitutions increased the amount of NA reaching the cell surface; activity per enzyme was unaffected by NA-V234M and slightly decreased by NA-R222Q [10]. As a result, in 2008 after wide-spread circulation of viruses carrying permissive secondary antiviral substitutions the NA-H275Y could establish circulation as a stable lineage and spread rapidly and unexpectedly around the world without any therapeutic selection pressure.

In 2009 the pandemic A(H1N1)pdm09 viruses replaced the previously oseltamivir-resistant seasonal A(H1N1) viruses and have been circulating worldwide as seasonal viruses ever since [26]. These 2009 pandemic A(H1N1) viruses remain largely oseltamivir-sensitive due to a reassortment event [11]. The introduction of NA-H275Y in A/California/4/2009 NA also resulted in a sharp decrease in total surface-expressed activity in these A(H1N1)pdm09 viruses as in the former seasonal A(H1N1) viruses [9]. However, the evolutionary development of the A(H1N1)pdm09 viruses, led to the establishment of the secondary NA substitutions NA-V241I and NA-N369K. In Germany the prevalence of viruses carrying these substitutions was about 60% in the 2010-2011 season. Since influenza season 2011-2012 all A(H1N1)pdm09 viruses analyzed showed the valine-isoleucine substitution at NA-241 and the aspartate acid were substituted by leucine at NA-369 position.

Using reverse-engineered viruses, these two substitutions, NA-V241I and NA-N369K, were shown to confer robust fitness to A(H1N1)pdm09 viruses carrying the NA-H275Y by increasing surface expression and enzymatic activity of the enzyme [17]. Nevertheless, the prevalence of resistant viruses in Germany and worldwide remained low (<1%) [15, 27-29]. In contrast to previous seasonal viruses, these permissive secondary neuraminidase substitutions did not lead to the emergence and rapid spread of resistant viruses.

These permissive secondary antiviral mutations have so far not supported the emergence of resistant lineages. A possible reason for this could be the structure of pandemic N1pdm09 NA, which generally differs from that of group N1 and group N2 neuraminidases. For example, the NA of H1N1pdm09 lacks the 150-cavity that is typical for group 1 NA, and additionally the salt bridge between Asp-147 and His-150 that is typical for group 2 NA [30, 31]. These structural features may require several permissive mutations to occur in the NA in a series of gradual steps to support the emergence of a resistant variant and establish it as a stable lineage.

There are indications of two further such substitutions, namely NA-I223V and NA-S247N. Our data show that the NA substitution from isoleucine to valine at position 223 and the change of serine by asparagine at position 247 have only a marginal effect on the susceptibility of A(H1N1)pdm09 viruses to NA inhibitors. However, NAs bearing both substitutions demonstrated a more than 10-fold reduction in susceptibility to oseltamivir and almost a 4-fold reduction in susceptibility to zanamivir compared to wild type virus [32]. Enzymatic analysis has proven that the NA-I223V substitution can offset the loss of affinity of NA for its substrate resulting from the NA-H275Y resistance substitution, while simultaneously boosting resistance [33]. Thermodynamic and structural analyses showed that the combination of NA-H275Y with NA-I223V or NA-S247N leads to an extreme reduction in the inhibitory potential of oseltamivir [19].

After a period of quiescence during the COVID-19 pandemic, the incidence of influenza has increased following the relaxation of non-pharmaceutical measures [4]. In Germany, A(H1N1)pdm09 viruses circulated with very low prevalence in the 2021-2022 and 2022-2023 influenza seasons, while the 2023-2024 season was dominated by A(H1N1)pdm09 (in preparation). In this season, a very small increase in the secondary substitutions NA-I223V and NA-S247N was observed compared to the pre-Covid-19 pandemic influenza seasons. The recently analysis of collected sequence data available through GISAID indicate an increased prevalence of these two permissive substitutions in globally circulating A(H1N1)pdm09 viruses with highest incidences of NA-I223V in August and October 2023 and of NA-S247N in September [32]. During these months, influenza viruses circulate primarily in the Southern Hemisphere and are considered precursors to those circulating in the northern hemisphere during the winter (October-April). A stronger spread of A(H1N1)pdm09 viruses with permissive secondary mutations could therefore be expected for the coming influenza seasons. Based on the experience with the unexpected emergence and subsequent strong spread of oseltamivir-resistant earlier seasonal A(H1N1) viruses, it is necessary to closely monitor the evolution of pandemic A(H1N1)pdm09 viruses. It seems likely that the viruses have reached the next stage in the evolution of prerequisite viruses that enable the emergence and spread of stable lineages of resistant viruses, in which the substitutions NA-I223V and NA-S247N may have been added in 2023-2024 after the appearance of the two permissive substitutions NA-V241I and NA-N369K in 2011. Continued monitoring of these permissive mutations is essential to ensure preparedness for the potential emergence of neuraminidase-resistant viruses. In parallel, the advancement of alternative antiviral agents, including polymerase inhibitors, is of great urgency.

## 5. Conclusions

Although a relationship between permissive secondary NA mutations and the emergence and spread of oseltamivir-resistant influenza viruses has been described in the past, little is known about the importance of permissive mutations in molecular evolution. In the absence of treatment selection pressure, permissive secondary mutations could support selection for resistant variants by compensating for their reduced viral fitness. As a result, the spread of antiviral resistant influenza viruses throughout the community seems likely. It is therefore important to closely monitor the evolution of circulating influenza viruses. Our data showed that the prevalence of permissive secondary antiviral mutations in the A(H1N1)pdm09 neuraminidase increased during the last influenza season.

## Supplementary Materials

The following supporting information can be downloaded at: www.mdpi.com/xxx/s1, Table S1: Accession number (Gisaid) of influenza A(H1N1)pdm09 viruses isolated in Germany between 2009 and 2024

## Author Contributions

Conceptualization, S.D. R.D. and B.S.; methodology, S.D., MW, JM; validation, S.D., J.M. and M.W.; formal analysis, S.D.; investigation, S.D., J.M., M.W., A.H.; data curation, A.H., J.M.; writing—original draft preparation, S.D.; writing—review and editing, S.D., B.S., R.D., A.H.; visualization, M.W., A.H.; supervision, B.S. and R.D.; project administration, S.D. All authors have read and agreed to the published version of the manuscript.

## Institutional Review Board Statement

The study was conducted in accordance with the Declaration of Helsinki, and approved by the Charité-Universitätsmedizin Berlin Ethical Board (reference EA2/126/11) and sentinel surveillance is covered by German legislation (§13, §14, Protection against Infection Act).

## Informed Consent Statement

Informed consent was obtained from all subjects involved in the study.

## Data Availability Statement

The original data generated as genome sequences presented in the study are openly available in Gisaid database at Gisaid.org. Generated IC_50_s for respective virus isolates are listed in the manuscript or are available on request from the corresponding author due to data protection rules.

## Acknowledgments

We extent our gratitude to the patients, their families and the sentinel clinicians who have made this study possible. We would like to thank the team of the German NIC in particular S.Tietze, M. Sohn, S. Hafemann, K. Merkel, U. Hopf-Guevara, K. Madaj, M. Smallfield, H. Fischer, M. Adam, B. Mischke, A. Schindel, J. Tesch, and S. Muschter for technical assistance; Ute Preuß, Anabel Hales, Louisa Schmidt and Mariella Szafraniec for outstanding support in recruiting sentinel physicians and coordinating the liaison with them; and Bernd Reinhardt for providing invaluable data management support. We would like to thank Andrea Thürmer and Aleksandar Radonic for the next generation sequencing of the influenza genomes and Katja Winter and Sandra Kaiser for the bioinformatic analysis of the NGS data.

## Disclaimer/Publisher’s Note

The statements, opinions and data contained in all publications are solely those of the individual author(s) and contributor(s) and not of MDPI and/or the editor(s). MDPI and/or the editor(s) disclaim responsibility for any injury to people or property resulting from any ideas, methods, instructions or products referred to in the content.

