## Supplementary material for "Increase of permissive secondary antiviral mutations in the evolution of A(H1N1)pdm09 influenza virus neuraminidases": Table S1

Table 1S. Accession number (Gisaid) of influenza A(H1N1)pdm09 viruses isolated in Germany between 2009 and 2024

| | | | |
|---|---|---|---|
| EPI_ISL_73769 | A/Sachsen-Anhalt/138/2009 | EPI_ISL_19140819 | A/Baden-Wuerttemberg/4/2010 |
| EPI_ISL_73768 | A/Sachsen-Anhalt/137/2009 | EPI_ISL_19140820 | A/Baden-Wuerttemberg/8/2010 |
| EPI_ISL_73766 | A/Sachsen-Anhalt/134/2009 | EPI_ISL_19140821 | A/Baden-Wuerttemberg/12/2009 |
| EPI_ISL_73764 | A/Sachsen-Anhalt/121/2009 | EPI_ISL_19140822 | A/Baden-Wuerttemberg/10/2009 |
| EPI_ISL_73763 | A/Sachsen-Anhalt/117/2009 | EPI_ISL_19140823 | A/Baden-Wuerttemberg/10/2010 |
| EPI_ISL_73760 | A/Sachsen/180/2009 | EPI_ISL_19140824 | A/Baden-Wuerttemberg/11/2009 |
| EPI_ISL_73759 | A/Sachsen/156/2009 | EPI_ISL_19140825 | A/Bayern/73/2009 |
| EPI_ISL_73750 | A/Niedersachsen/92/2009 | EPI_ISL_19140826 | A/Bayern/74/2009 |
| EPI_ISL_73741 | A/Niedersachsen/336/2009 | EPI_ISL_19140827 | A/Bayern/63/2009 |
| EPI_ISL_73739 | A/Niedersachsen/216/2009 | EPI_ISL_19140828 | A/Bayern/69/2009 |
| EPI_ISL_73738 | A/Niedersachsen/188/2009 | EPI_ISL_19140829 | A/Bayern/77/2009 |
| EPI_ISL_73737 | A/Niedersachsen/134/2009 | EPI_ISL_19140830 | A/Bayern/78/2009 |
| EPI_ISL_73736 | A/Niedersachsen/109/2009 | EPI_ISL_19140831 | A/Bayern/84/2009 |
| EPI_ISL_73735 | A/Niedersachsen/104/2009 | EPI_ISL_19140832 | A/Bayern/86/2009 |
| EPI_ISL_73689 | A/Bayern/79/2009 | EPI_ISL_19140833 | A/Bayern/96/2009 |
| EPI_ISL_73710 | A/Berlin/97/2009 | EPI_ISL_19140834 | A/Bayern/107/2009 |
| EPI_ISL_73698 | A/Brandenburg/43/2009 | EPI_ISL_19140835 | A/Bayern/150/2009 |
| EPI_ISL_73742 | A/Niedersachsen/343/2009 | EPI_ISL_19140836 | A/Bayern/154/2009 |
| EPI_ISL_73743 | A/Niedersachsen/352/2009 | EPI_ISL_19140837 | A/Bayern/125/2009 |
| EPI_ISL_19140806 | A/Schleswig-Holstein/16/2009 | EPI_ISL_19140838 | A/Bayern/128/2009 |
| EPI_ISL_19140807 | A/Bayern/75/2009 | EPI_ISL_19140839 | A/Bayern/117/2009 |
| EPI_ISL_19140808 | A/Hessen/47/2009 | EPI_ISL_19140840 | A/Bayern/9/2010 |
| EPI_ISL_19140809 | A/Nordrhein-Westfalen/74/2009 | EPI_ISL_19140814 | A/Baden-Wuerttemberg/487/2009 |
| EPI_ISL_19140810 | A/Hessen/48/2009 | EPI_ISL_19140815 | A/Baden-Wuerttemberg/487/2009 |
| EPI_ISL_19140811 | A/Baden-Wuerttemberg/459/2009 | EPI_ISL_19140816 | A/Baden-Wuerttemberg/490/2009 |
| EPI_ISL_19140812 | A/Baden-Wuerttemberg/462/2009 | EPI_ISL_19140817 | A/Baden-Wuerttemberg/502/2009 |
| EPI_ISL_19140813 | A/Baden-Wuerttemberg/464/2009 | EPI_ISL_19140818 | A/Baden-Wuerttemberg/511/2009 |
| EPI_ISL_19140841 | A/Berlin/74/2009 | EPI_ISL_19140873 | A/Hessen/10/2009 |
| EPI_ISL_19140842 | A/Berlin/72/2009 | EPI_ISL_19140874 | A/Mecklenburg-Vorpommern/1/2009 |
| EPI_ISL_19140843 | A/Berlin/80/2009 | EPI_ISL_19140875 | A/Mecklenburg-Vorpommern/6/2009 |
| EPI_ISL_19140844 | A/Berlin/78/2009 | EPI_ISL_19140876 | A/Mecklenburg-Vorpommern/10/09 |
| EPI_ISL_19140845 | A/Berlin/79/2009 | EPI_ISL_19140877 | A/Mecklenburg-Vorpommern/11/2009 |

Table 1S. Accession number (Gisaid) of influenza A(H1N1)pdm09 viruses isolated in Germany between 2009 and 2024

| | | | |
|---|---|---|---|
| EPI_ISL_19140846 | A/Berlin/75/2009 | EPI_ISL_19140878 | A/Mecklenburg-Vorpommern/10/2009 |
| EPI_ISL_19140847 | A/Berlin/87/2009 | EPI_ISL_19140879 | A/Niedersachsen/339/2009 |
| EPI_ISL_19140848 | A/Berlin/88/2009 | EPI_ISL_19140880 | A/Niedersachsen/84/2009 |
| EPI_ISL_19140849 | A/Berlin/90/2009 | EPI_ISL_19140881 | A/Niedersachsen/87/2009 |
| EPI_ISL_19140850 | A/Berlin/83/2009 | EPI_ISL_19140882 | A/Niedersachsen/88/2009 |
| EPI_ISL_19140851 | A/Berlin/121/2009 | EPI_ISL_19140883 | A/Niedersachsen/415/2009 |
| EPI_ISL_19140852 | A/Berlin/130/2009 | EPI_ISL_19140884 | A/Niedersachsen/10/2010 |
| EPI_ISL_19140853 | A/Berlin/147/2009 | EPI_ISL_19140885 | A/Nordrhein-Westfalen/100/2009 |
| EPI_ISL_19140854 | A/Berlin/164/2009 | EPI_ISL_19140886 | A/Nordrhein-Westfalen/87/2009 |
| EPI_ISL_19140855 | A/Berlin/185/2009 | EPI_ISL_19140887 | A/Nordrhein-Westfalen/94/2009 |
| EPI_ISL_19140856 | A/Berlin/187/2009 | EPI_ISL_19140888 | A/Nordrhein-Westfalen/99/2009 |
| EPI_ISL_19140857 | A/Berlin/210/2009 | EPI_ISL_19140889 | A/Nordrhein-Westfalen/103/2009 |
| EPI_ISL_19140858 | A/Berlin/10/2009 | EPI_ISL_19140890 | A/Nordrhein-Westfalen/106/2009 |
| EPI_ISL_19140859 | A/Berlin/10/2010 | EPI_ISL_19140891 | A/Nordrhein-Westfalen/113/2009 |
| EPI_ISL_19140860 | A/Berlin/143/2009 | EPI_ISL_19140892 | A/Nordrhein-Westfalen/137/2009 |
| EPI_ISL_19140861 | A/Brandenburg/38/2009 | EPI_ISL_19140893 | A/Nordrhein-Westfalen/152/2009 |
| EPI_ISL_19140862 | A/Brandenburg/28/2009 | EPI_ISL_19140894 | A/Nordrhein-Westfalen/184/2009 |
| EPI_ISL_19140863 | A/Brandenburg/37/2009 | EPI_ISL_19140895 | A/Nordrhein-Westfalen/163/2009 |
| EPI_ISL_19140864 | A/Brandenburg/42/2009 | EPI_ISL_19140896 | A/Nordrhein-Westfalen/201/2009 |
| EPI_ISL_19140865 | A/Brandenburg/24/2009 | EPI_ISL_19140897 | A/Rheinland-Pfalz/43/2009 |
| EPI_ISL_19140866 | A/Brandenburg/34/2009 | EPI_ISL_19140898 | A/Rheinland-Pfalz/41/2009 |
| EPI_ISL_19140867 | A/Brandenburg/10/2009 | EPI_ISL_19140899 | A/Rheinland-Pfalz/44/2009 |
| EPI_ISL_19140868 | A/Bremen/18/2009 | EPI_ISL_19140900 | A/Rheinland-Pfalz/81/2009 |
| EPI_ISL_19140869 | A/Bremen/20/2009 | EPI_ISL_19140901 | A/Rheinland-Pfalz/86/2009 |
| EPI_ISL_19140870 | A/Hamburg/6/2009 | EPI_ISL_19140902 | A/Rheinland-Pfalz/10/2009 |
| EPI_ISL_19140871 | A/Hamburg/14/2009 | EPI_ISL_19140903 | A/Rheinland-Pfalz/11/2009 |
| EPI_ISL_19140872 | A/Hessen/61/2009 | EPI_ISL_19140904 | A/Saarland/3/2009 |
| EPI_ISL_19140905 | A/Saarland/16/2009 | EPI_ISL_19140936 | A/Nordrhein-Westfalen/173/2009 |
| EPI_ISL_19140906 | A/Sachsen/96/2009 | EPI_ISL_19140937 | A/Nordrhein-Westfalen/174/2009 |
| EPI_ISL_19140907 | A/Sachsen/93/2009 | EPI_ISL_19140938 | A/Nordrhein-Westfalen/175/2009 |
| EPI_ISL_19140908 | A/Sachsen/92/2009 | EPI_ISL_19140939 | A/Niedersachsen/850/2009 |
| EPI_ISL_19140909 | A/Sachsen/10/2009 | EPI_ISL_19140940 | A/Niedersachsen/866/2009 |

Table 1S. Accession number (Gisaid) of influenza A(H1N1)pdm09 viruses isolated in Germany between 2009 and 2024

| | | | |
|---|---|---|---|
| EPI_ISL_19140910 | A/Sachsen/97/2009 | EPI_ISL_19140941 | A/Sachsen/183/2009 |
| EPI_ISL_19140911 | A/Sachsen/129/2009 | EPI_ISL_19140942 | A/Sachsen-Anhalt/178/2009 |
| EPI_ISL_19140912 | A/Sachsen/154/2009 | EPI_ISL_19141140 | *A/Berlin/12/2010* |
| EPI_ISL_19140913 | A/Sachsen/194/2009 | EPI_ISL_19141141 | *A/Berlin/14/2010* |
| EPI_ISL_19140969 | A/Sachsen/194/2009 Res | EPI_ISL_19141142 | *A/Berlin/15/2010* |
| EPI_ISL_19140914 | A/Sachsen/4/2010 | EPI_ISL_19141143 | *A/Berlin/17/2010* |
| EPI_ISL_19140915 | A/Sachsen-Anhalt/9/2010 | EPI_ISL_19141144 | *A/Berlin/18/2010* |
| EPI_ISL_19140916 | A/Schleswig-Holstein/13/2009 | EPI_ISL_19141189 | A/Baden-Wuerttemberg/13/2010 |
| EPI_ISL_19140917 | A/Thueringen/173/2009 | EPI_ISL_19141190 | A/Baden-Wuerttemberg/14/2010 |
| EPI_ISL_19140918 | A/Thueringen/176/2009 | EPI_ISL_19141191 | A/Nordrhein-Westfalen/11/2010 |
| EPI_ISL_19140919 | A/Thueringen/189/2009 | EPI_ISL_19141145 | A/Bayern/4/2011 |
| EPI_ISL_19140920 | A/Thueringen/201/2009 | EPI_ISL_19141146 | A/Bayern/7/2011 |
| EPI_ISL_19140921 | A/Thueringen/227/2009 | EPI_ISL_19141147 | A/Berlin/14/2011 |
| EPI_ISL_19140922 | A/Sachsen-Anhalt/127/2009 | EPI_ISL_19141148 | A/Bayern/39/2011 |
| EPI_ISL_19140923 | A/Sachsen-Anhalt/135/2009 | EPI_ISL_19141149 | A/Brandenburg/1/2011 |
| EPI_ISL_19140924 | A/Niedersachsen/330/2009 | EPI_ISL_19141150 | A/Berlin/6/2011 |
| EPI_ISL_19140925 | A/Sachsen-Anhalt/144/2009 | EPI_ISL_19141151 | A/Berlin/8/2011 |
| EPI_ISL_19140926 | A/Niedersachsen/115/2009 | EPI_ISL_19141152 | A/Berlin/23/2011 |
| EPI_ISL_19140927 | A/Sachsen-Anhalt/125/2009 | EPI_ISL_19141153 | A/Berlin/38/2011 |
| EPI_ISL_19140928 | A/Sachsen-Anhalt/119/2009 | EPI_ISL_19141154 | A/Bremen/3/2011 |
| EPI_ISL_19140929 | A/Sachsen-Anhalt/122/2009 | EPI_ISL_19141155 | A/Hamburg/3/2011 |
| EPI_ISL_19140930 | A/Niedersachsen/337/2009 | EPI_ISL_19141156 | A/Hessen/3/2011 |
| EPI_ISL_19140931 | A/Niedersachsen/338/2009 | EPI_ISL_19141192 | A/Nordrhein-Westfalen/5/2011 |
| EPI_ISL_19140932 | A/Sachsen-Anhalt/124/2009 | EPI_ISL_19141193 | A/Nordrhein-Westfalen/6/2011 |
| EPI_ISL_19140933 | A/Niedersachsen/91/2009 | EPI_ISL_19141194 | A/Nordrhein-Westfalen/16/2011 |
| EPI_ISL_19140934 | A/Nordrhein-Westfalen/171/2009 | EPI_ISL_19141195 | A/Nordrhein-Westfalen/53/2011 |
| EPI_ISL_19140935 | A/Nordrhein-Westfalen/172/2009 | EPI_ISL_19141157 | A/Niedersachsen/7/2011 |
| EPI_ISL_19141158 | A/Niedersachsen/15/2011 | EPI_ISL_19141200 | A/Nordrhein-Westfalen/14/2010 |
| EPI_ISL_19141159 | A/Niedersachsen/16/2011 | EPI_ISL_19141183 | A/Niedersachsen/11/2011 |
| EPI_ISL_19141160 | A/Niedersachsen/80/2011 | EPI_ISL_19141184 | A/Niedersachsen/13/2010 |
| EPI_ISL_19141161 | A/Rheinland-Pfalz/5/2011 | EPI_ISL_19141185 | A/Niedersachsen/11/2010 |
| EPI_ISL_19141162 | A/Rheinland-Pfalz/7/2011 | EPI_ISL_19141186 | A/Niedersachsen/12/2010 |

Table 1S. Accession number (Gisaid) of influenza A(H1N1)pdm09 viruses isolated in Germany between 2009 and 2024

| | | | |
|---|---|---|---|
| EPI_ISL_19141163 | A/Rheinland-Pfalz/42/2011 | EPI_ISL_19141187 | A/Sachsen/28/2011 |
| EPI_ISL_19141164 | A/Rheinland-Pfalz/55/2011 | EPI_ISL_19141188 | A/Sachsen/10/2011 |
| EPI_ISL_19141165 | A/Saarland/1/2011 | EPI_ISL_19141201 | A/Rheinland-Pfalz/17/2011 |
| EPI_ISL_19141166 | A/Sachsen/2/2011 | EPI_ISL_19141214 | A/Baden-Wuerttemberg/60/2012 |
| EPI_ISL_19141167 | A/Sachsen/8/2011 | EPI_ISL_19141215 | A/Bayern/28/2012 |
| EPI_ISL_19141168 | A/Schleswig-Holstein/3/2011 | EPI_ISL_19141216 | A/Niedersachsen/10/2012 |
| EPI_ISL_19141169 | A/Thueringen/2/2011 | EPI_ISL_19141217 | A/Berlin/136/2012 |
| EPI_ISL_19141170 | A/Bayern/10/2010 | EPI_ISL_19141218 | A/Baden-Wuerttemberg/172/2012 |
| EPI_ISL_19141171 | A/Bayern/11/2010 | EPI_ISL_19141219 | A/Bayern/86/2012 |
| EPI_ISL_19141172 | A/Berlin/13/2011 | EPI_ISL_19141220 | A/Baden-Wuerttemberg/28/2012 |
| EPI_ISL_19141173 | A/Berlin/10/2011 | EPI_ISL_19141221 | A/Baden-Wuerttemberg/109/2012 |
| EPI_ISL_19141174 | A/Berlin/11/2010 | EPI_ISL_19141222 | A/Baden-Wuerttemberg/149/2012 |
| EPI_ISL_19141142 | A/Berlin/15/2010 | EPI_ISL_19141223 | A/Niedersachsen/6/2012 |
| EPI_ISL_19141140 | A/Berlin/12/2010 | EPI_ISL_19141224 | A/Baden-Wuerttemberg/125/2012 |
| EPI_ISL_19141175 | A/Berlin/11/2011 | EPI_ISL_19141438 | A/Bayern/91/2012 |
| EPI_ISL_19141176 | A/Berlin/12/2011 | EPI_ISL_19141439 | A/Berlin/167/2012 |
| EPI_ISL_19141177 | A/Berlin/13/2010 | EPI_ISL_19141440 | A/Nordrhein-Westfalen/38/2012 |
| EPI_ISL_19141141 | A/Berlin/14/2010 | EPI_ISL_19141441 | A/Baden-Wuerttemberg/80/2013 |
| EPI_ISL_19141178 | A/Berlin/16/2010 | EPI_ISL_19141442 | A/Hessen/4/2013 |
| EPI_ISL_19141179 | A/Bremen/2/2010 | EPI_ISL_19141443 | A/Rheinland-Pfalz/2/2013 |
| EPI_ISL_19141180 | A/Bremen/3/2010 | EPI_ISL_19141444 | A/Sachsen/6/2013 |
| EPI_ISL_19141196 | A/Baden-Wuerttemberg/10/2011 | EPI_ISL_19141445 | A/Bayern/93/2012 |
| EPI_ISL_19141197 | A/Baden-Wuerttemberg/11/2011 | EPI_ISL_19141446 | A/Thueringen/42/2012 |
| EPI_ISL_19141198 | A/Baden-Wuerttemberg/12/2011 | EPI_ISL_19151069 | A/Brandenburg/1/2014 |
| EPI_ISL_19141199 | A/Baden-Wuerttemberg/11/2010 | EPI_ISL_19151057 | A/Berlin/6/2014 |
| EPI_ISL_19141181 | A/Hessen/11/2011 | EPI_ISL_19151058 | A/Berlin/30/2014 |
| EPI_ISL_19141182 | A/Hessen/10/2011 | EPI_ISL_19151059 | A/Schleswig-Holstein/2/2014 |
| EPI_ISL_19151060 | A/Berlin/54/2014 | EPI_ISL_19151093 | A/Niedersachsen/68/2015 |
| EPI_ISL_19151061 | A/Berlin/55/2014 | EPI_ISL_19151094 | A/Niedersachsen/72/2015 |
| EPI_ISL_19151062 | A/Berlin/40/2014 | EPI_ISL_19151095 | A/Rheinland-Pfalz/53/2015 |
| EPI_ISL_19151063 | A/Berlin/10/2014 | EPI_ISL_19151096 | A/Rheinland-Pfalz/54/2015 |
| EPI_ISL_19151064 | A/Bayern/12/2014 | EPI_ISL_19151097 | A/Rheinland-Pfalz/10/2015 |

Table 1S. Accession number (Gisaid) of influenza A(H1N1)pdm09 viruses isolated in Germany between 2009 and 2024

| | | | |
|---|---|---|---|
| EPI_ISL_19151065 | A/Sachsen/10/14 | EPI_ISL_19151098 | A/Thueringen/43/2015 |
| EPI_ISL_19151066 | A/Sachsen/10/2014 | EPI_ISL_202714 | A/Bayern/146/2015 |
| EPI_ISL_19151070 | A/Baden-Wuerttemberg/10/2014 | EPI_ISL_208323 | A/Bayern/148/2015 |
| EPI_ISL_19151067 | A/Berlin/13/2014 | EPI_ISL_203628 | A/Bayern/149/2015 |
| EPI_ISL_19151068 | A/Brandenburg/10/2014 | EPI_ISL_208333 | A/Bayern/151/2015 |
| EPI_ISL_19151071 | A/Berlin/11/2014 | EPI_ISL_202562 | A/Berlin/166/2015 |
| EPI_ISL_19151072 | A/Bremen/8/2014 | EPI_ISL_202715 | A/Berlin/167/2015 |
| EPI_ISL_19151073 | A/Berlin/12/2014 | EPI_ISL_203623 | A/Berlin/170/2015 |
| EPI_ISL_19151074 | A/Berlin/81/2014 | EPI_ISL_203624 | A/Berlin/171/2015 |
| EPI_ISL_19151075 | A/Hamburg/7/2014 | EPI_ISL_208355 | A/Berlin/174/2015 |
| EPI_ISL_19151076 | A/Rheinland-Pfalz/12/2014 | EPI_ISL_208327 | A/Berlin/175/2015 |
| EPI_ISL_19151077 | A/Rheinland-Pfalz/14/2014 | EPI_ISL_208322 | A/Baden-Wuerttemberg/253/2015 |
| EPI_ISL_19151078 | A/Rheinland-Pfalz/15/2014 | EPI_ISL_208324 | A/Baden-Wuerttemberg/254/2015 |
| EPI_ISL_19151079 | A/Schleswig-Holstein/9/2014 | EPI_ISL_208330 | A/Berlin/176/2015 |
| EPI_ISL_19151080 | A/Bayern/110/2015 | EPI_ISL_202564 | A/Hamburg/24/2015 |
| EPI_ISL_19151081 | A/Brandenburg/29/2015 | EPI_ISL_202716 | A/Hamburg/25/2015 |
| EPI_ISL_19151082 | A/Berlin/78/2015 | EPI_ISL_202565 | A/Hamburg/26/2015 |
| EPI_ISL_19151083 | A/Berlin/113/2015 | EPI_ISL_208329 | A/Mecklenburg-Vorpommern/20/2015 |
| EPI_ISL_19151084 | A/Berlin/97/2015 | EPI_ISL_208336 | A/Berlin/177/2015 |
| EPI_ISL_19151085 | A/Bremen/23/2015 | EPI_ISL_208326 | A/Mecklenburg-Vorpommern/21/2015 |
| EPI_ISL_19151086 | A/Baden-Wuerttemberg/14/2015 | EPI_ISL_202717 | A/Nordrhein-Westfalen/102/2015 |
| EPI_ISL_19151087 | A/Hamburg/23/2015 | EPI_ISL_202718 | A/Nordrhein-Westfalen/103/2015 |
| EPI_ISL_19151088 | A/Hamburg/10/2015 | EPI_ISL_203629 | A/Nordrhein-Westfalen/105/2015 |
| EPI_ISL_19151089 | A/Hessen/37/2015 | EPI_ISL_208337 | A/Nordrhein-Westfalen/106/2015 |
| EPI_ISL_19151090 | A/Nordrhein-Westfalen/77/2015 | EPI_ISL_202563 | A/Bremen/25/2015 |
| EPI_ISL_19151091 | A/Nordrhein-Westfalen/83/2015 | EPI_ISL_203626 | A/Rheinland-Pfalz/55/2015 |
| EPI_ISL_19151092 | A/Nordrhein-Westfalen/93/2015 | EPI_ISL_212442 | A/Sachsen/102/2015 |
| EPI_ISL_208328 | A/Sachsen/104/2015 | EPI_ISL_222727 | A/Bayern/16/2016 |
| EPI_ISL_203625 | A/Sachsen-Anhalt/91/2015 | EPI_ISL_222729 | A/Bayern/21/2016 |
| EPI_ISL_208332 | A/Thueringen/101/2015 | EPI_ISL_222731 | A/Bayern/23/2016 |
| EPI_ISL_203627 | A/Bremen/26/2015 | EPI_ISL_222736 | A/Bayern/24/2016 |
| EPI_ISL_208338 | A/Hamburg/1/2016 | EPI_ISL_213449 | A/Bayern/30/2016 |

Table 1S. Accession number (Gisaid) of influenza A(H1N1)pdm09 viruses isolated in Germany between 2009 and 2024

| | | | |
|---|---|---|---|
| EPI_ISL_208334 | A/Thueringen/1/2016 | EPI_ISL_222613 | A/Bayern/36/2016 |
| EPI_ISL_208335 | A/Thueringen/2/2016 | EPI_ISL_222620 | A/Bayern/42/2016 |
| EPI_ISL_208325 | A/Bremen/28/2015 | EPI_ISL_222655 | A/Bayern/45/2016 |
| EPI_ISL_19159374 | A/Bremen/27/2015 | EPI_ISL_222631 | A/Bayern/47/2016 |
| EPI_ISL_19159375 | A/Thueringen/103/2015 | EPI_ISL_223783 | A/Bayern/50/2016 |
| EPI_ISL_19159376 | A/Bayern/94/2016 | EPI_ISL_223789 | A/Bayern/52/2016 |
| EPI_ISL_19159377 | A/Bayern/20/2016 | EPI_ISL_222724 | A/Bayern/54/2016 |
| EPI_ISL_19159378 | A/Berlin/1/2016 | EPI_ISL_222725 | A/Bayern/56/2016 |
| EPI_ISL_19159379 | A/Berlin/3/2016 | EPI_ISL_222738 | A/Bayern/59/2016 |
| EPI_ISL_19159380 | A/Rheinland-Pfalz/1/2016 | EPI_ISL_222619 | A/Bayern/65/2016 |
| EPI_ISL_19159381 | A/Schleswig-Holstein/4/2016 | EPI_ISL_222633 | A/Bayern/66/2016 |
| EPI_ISL_19159386 | A/Bremen/10/2015 | EPI_ISL_222646 | A/Bayern/67/2016 |
| EPI_ISL_19159382 | A/Berlin/103/2016 | EPI_ISL_222649 | A/Bayern/68/2016 |
| EPI_ISL_19159383 | A/Berlin/168/2015 | EPI_ISL_211850 | A/Brandenburg/2/2016 |
| EPI_ISL_19159384 | A/Berlin/169/2015 | EPI_ISL_211855 | A/Brandenburg/3/2016 |
| EPI_ISL_19159385 | A/Berlin/172/2015 | EPI_ISL_212497 | A/Brandenburg/9/2016 |
| EPI_ISL_19159684 | A/Bayern/6/2016 | EPI_ISL_213308 | A/Brandenburg/13/2016 |
| EPI_ISL_19159381 | A/Schleswig-Holstein/3/2016 | EPI_ISL_222605 | A/Brandenburg/15/2016 |
| EPI_ISL_19159386 | A/Nordrhein-Westfalen/2/2016 | EPI_ISL_213448 | A/Brandenburg/17/2016 |
| EPI_ISL_19159382 | A/Thueringen/105/2015 | EPI_ISL_222733 | A/Brandenburg/18/2016 |
| EPI_ISL_222723 | A/Thueringen/106/2015 | EPI_ISL_223786 | A/Brandenburg/22/2016 |
| EPI_ISL_208339 | A/Bayern/1/2016 | EPI_ISL_208344 | A/Berlin/1/2016 |
| EPI_ISL_208347 | A/Bayern/2/2016 | EPI_ISL_208342 | A/Berlin/2/2016 |
| EPI_ISL_208345 | A/Bayern/3/2016 | EPI_ISL_211848 | A/Berlin/4/2016 |
| EPI_ISL_211853 | A/Bayern/5/2016 | EPI_ISL_208349 | A/Berlin/5/2016 |
| EPI_ISL_2120501 | A/Bayern/7/2016 | EPI_ISL_211854 | A/Berlin/6/2016 |
| EPI_ISL_213442 | A/Bayern/12/2016 | EPI_ISL_208346 | A/Berlin/9/2016 |
| EPI_ISL_212495 | A/Berlin/13/2016 | EPI_ISL_222614 | A/Baden-Wuerttemberg/39/2016 |
| EPI_ISL_212498 | A/Berlin/15/2016 | EPI_ISL_222621 | A/Baden-Wuerttemberg/52/2016 |
| EPI_ISL_212503 | A/Berlin/25/2016 | EPI_ISL_222623 | A/Baden-Wuerttemberg/53/2016 |
| EPI_ISL_213437 | A/Berlin/30/2016 | EPI_ISL_222622 | A/Baden-Wuerttemberg/56/2016 |
| EPI_ISL_222609 | A/Berlin/35/2016 | EPI_ISL_222632 | A/Baden-Wuerttemberg/61/2016 |

Table 1S. Accession number (Gisaid) of influenza A(H1N1)pdm09 viruses isolated in Germany between 2009 and 2024

| | | | |
|---|---|---|---|
| EPI_ISL_213444 | A/Berlin/38/2016 | EPI_ISL_222647 | A/Baden-Wuerttemberg/72/2016 |
| EPI_ISL_222611 | A/Berlin/39/2016 | EPI_ISL_223782 | A/Baden-Wuerttemberg/84/2016 |
| EPI_ISL_213758 | A/Berlin/40/2016 | EPI_ISL_211851 | A/Hamburg/2/2016 |
| EPI_ISL_213445 | A/Berlin/41/2016 | EPI_ISL_222599 | A/Hamburg/3/2016 |
| EPI_ISL_222608 | A/Berlin/45/2016 | EPI_ISL_212504 | A/Hamburg/6/2016 |
| EPI_ISL_218495 | A/Berlin/55/2016 | EPI_ISL_213438 | A/Hessen/3/2016 |
| EPI_ISL_222730 | A/Berlin/60/2016 | EPI_ISL_222606 | A/Hessen/4/2016 |
| EPI_ISL_222616 | A/Berlin/71/2016 | EPI_ISL_222610 | A/Hessen/7/2016 |
| EPI_ISL_222624 | A/Berlin/76/2016 | EPI_ISL_218493 | A/Hessen/8/2016 |
| EPI_ISL_222638 | A/Berlin/77/2016 | EPI_ISL_218491 | A/Hessen/10/2016 |
| EPI_ISL_222726 | A/Berlin/84/2016 | EPI_ISL_222636 | A/Hessen/13/2016 |
| EPI_ISL_222648 | A/Berlin/85/2016 | EPI_ISL_222659 | A/Hessen/14/2016 |
| EPI_ISL_222650 | A/Berlin/86/2016 | EPI_ISL_222660 | A/Hessen/15/2016 |
| EPI_ISL_222651 | A/Berlin/87/2016 | EPI_ISL_208343 | A/Mecklenburg-Vorpommern/1/2016 |
| EPI_ISL_222607 | A/Bremen/1/2016 | EPI_ISL_222602 | A/Mecklenburg-Vorpommern/2/2016 |
| EPI_ISL_222737 | A/Bremen/5/2016 | EPI_ISL_212505 | A/Mecklenburg-Vorpommern/4/2016 |
| EPI_ISL_222735 | A/Bremen/6/2016 | EPI_ISL_222617 | A/Mecklenburg-Vorpommern/8/2016 |
| EPI_ISL_222628 | A/Bremen/8/2016 | EPI_ISL_208350 | A/Nordrhein-Westfalen/1/2016 |
| EPI_ISL_222640 | A/Bremen/9/2016 | EPI_ISL_211847 | A/Nordrhein-Westfalen/3/2016 |
| EPI_ISL_222639 | A/Bremen/10/2016 | EPI_ISL_212494 | A/Nordrhein-Westfalen/4/2016 |
| EPI_ISL_208341 | A/Baden-Wuerttemberg/1/2016 | EPI_ISL_222600 | A/Nordrhein-Westfalen/12/2016 |
| EPI_ISL_212506 | A/Baden-Wuerttemberg/7/2016 | EPI_ISL_213307 | A/Nordrhein-Westfalen/14/2016 |
| EPI_ISL_213447 | A/Baden-Wuerttemberg/16/2016 | EPI_ISL_222604 | A/Nordrhein-Westfalen/15/2016 |
| EPI_ISL_213446 | A/Baden-Wuerttemberg/17/2016 | EPI_ISL_213309 | A/Nordrhein-Westfalen/16/2016 |
| EPI_ISL_222734 | A/Baden-Wuerttemberg/25/2016 | EPI_ISL_218492 | A/Nordrhein-Westfalen/18/2016 |
| EPI_ISL_222612 | A/Baden-Wuerttemberg/34/2016 | EPI_ISL_212499 | A/Nordrhein-Westfalen/19/2016 |
| EPI_ISL_222741 | A/Baden-Wuerttemberg/37/2016 | EPI_ISL_222728 | A/Nordrhein-Westfalen/27/2016 |
| EPI_ISL_222618 | A/Nordrhein-Westfalen/34/2016 | EPI_ISL_222644 | A/Rheinland-Pfalz/35/2016 |
| EPI_ISL_223784 | A/Nordrhein-Westfalen/43/2016 | EPI_ISL_223788 | A/Rheinland-Pfalz/36/2016 |
| EPI_ISL_222625 | A/Nordrhein-Westfalen/57/2016 | EPI_ISL_223790 | A/Saarland/9/2016 |
| EPI_ISL_222601 | A/Niedersachsen/2/2016 | EPI_ISL_208352 | A/Sachsen/1/2016 |
| EPI_ISL_213441 | A/Niedersachsen/4/2016 | EPI_ISL_208353 | A/Sachsen/6/2016 |

Table 1S. Accession number (Gisaid) of influenza A(H1N1)pdm09 viruses isolated in Germany between 2009 and 2024

| | | | |
|---|---|---|---|
| EPI_ISL_218490 | A/Niedersachsen/8/2016 | EPI_ISL_211859 | A/Thueringen/104/2015 |
| EPI_ISL_213440 | A/Niedersachsen/9/2016 | EPI_ISL_212000 | A/Sachsen/14/2016 |
| EPI_ISL_218494 | A/Niedersachsen/12/2016 | EPI_ISL_222642 | A/Sachsen/54/2016 |
| EPI_ISL_213310 | A/Niedersachsen/25/2016 | EPI_ISL_222641 | A/Sachsen/61/2016 |
| EPI_ISL_222739 | A/Niedersachsen/29/2016 | EPI_ISL_223785 | A/Sachsen/66/2016 |
| EPI_ISL_222615 | A/Niedersachsen/35/2016 | EPI_ISL_223787 | A/Sachsen/68/2016 |
| EPI_ISL_222634 | A/Niedersachsen/37/2016 | EPI_ISL_222629 | A/Sachsen-Anhalt/76/2016 |
| EPI_ISL_222637 | A/Niedersachsen/38/2016 | EPI_ISL_222645 | A/Sachsen-Anhalt/62/2016 |
| EPI_ISL_222653 | A/Niedersachsen/39/2016 | EPI_ISL_212507 | A/Schleswig-Holstein/2/2016 |
| EPI_ISL_222643 | A/Niedersachsen/40/2016 | EPI_ISL_222627 | A/Schleswig-Holstein/16/2016 |
| EPI_ISL_222654 | A/Niedersachsen/41/2016 | EPI_ISL_208340 | A/Thueringen/3/2016 |
| EPI_ISL_222658 | A/Niedersachsen/42/2016 | EPI_ISL_211852 | A/Thueringen/4/2016 |
| EPI_ISL_222630 | A/Niedersachsen/54/2016 | EPI_ISL_211849 | A/Thueringen/5/2016 |
| EPI_ISL_222635 | A/Niedersachsen/55/2016 | EPI_ISL_213443 | A/Thueringen/14/2016 |
| EPI_ISL_222657 | A/Niedersachsen/56/2016 | EPI_ISL_222732 | A/Thueringen/18/2016 |
| EPI_ISL_211846 | A/Rheinland-Pfalz/2/2016 | EPI_ISL_211856 | A/Baden-Wuerttemberg/2/2016 |
| EPI_ISL_208351 | A/Rheinland-Pfalz/3/2016 | EPI_ISL_211860 | A/Sachsen/3/2016 |
| EPI_ISL_212493 | A/Rheinland-Pfalz/5/2016 | EPI_ISL_211861 | A/Sachsen/4/2016 |
| EPI_ISL_212496 | A/Rheinland-Pfalz/8/2016 | EPI_ISL_211862 | A/Sachsen/5/2016 |
| EPI_ISL_212500 | A/Rheinland-Pfalz/9/2016 | EPI_ISL_253536 | A/Nordrhein-Westfalen/26/2017 |
| EPI_ISL_212502 | A/Rheinland-Pfalz/11/2016 | EPI_ISL_253538 | A/Niedersachsen/31/2017 |
| EPI_ISL_213439 | A/Rheinland-Pfalz/12/2016 | EPI_ISL_253537 | A/Rheinland-Pfalz/38/2017 |
| EPI_ISL_213306 | A/Rheinland-Pfalz/13/2016 | EPI_ISL_253539 | A/Sachsen-Anhalt/3/2017 |
| EPI_ISL_222740 | A/Rheinland-Pfalz/25/2016 | EPI_ISL_19159932 | A/Baden-Wuerttemberg/256/2017 |
| EPI_ISL_222626 | A/Rheinland-Pfalz/32/2016 | EPI_ISL_19159930 | A/Baden-Wuerttemberg/16/2018 |
| EPI_ISL_222652 | A/Rheinland-Pfalz/33/2016 | EPI_ISL_19159931 | A/Hessen/45/2018 |
| EPI_ISL_222656 | A/Rheinland-Pfalz/34/2016 | EPI_ISL_301645 | A/Niedersachsen/17/2018 |
| EPI_ISL_19159933 | A/Saarland/25/2017 | EPI_ISL_301680 | A/Rheinland-Pfalz/19/2018 |
| EPI_ISL_288727 | A/Rheinland-Pfalz/50/2017 | EPI_ISL_301646 | A/Baden-Wuerttemberg/47/2018 |
| EPI_ISL_312788 | A/Bayern/48/2018 | EPI_ISL_311285 | A/Baden-Wuerttemberg/56/2018 |
| EPI_ISL_311284 | A/Nordrhein-Westfalen/19/2018 | EPI_ISL_301647 | A/Baden-Wuerttemberg/55/2018 |
| EPI_ISL_222603 | A/Sachsen/11/2016 | EPI_ISL_292227 | A/Nordrhein-Westfalen/135/2017 |

Table 1S. Accession number (Gisaid) of influenza A(H1N1)pdm09 viruses isolated in Germany between 2009 and 2024

| | | | |
|---|---|---|---|
| EPI_ISL_301644 | A/Bayern/7/2018 | EPI_ISL_301648 | A/Nordrhein-Westfalen/25/2018 |
| EPI_ISL_292228 | A/Bayern/96/2017 | EPI_ISL_301649 | A/Berlin/17/2018 |
| EPI_ISL_292125 | A/Saarland/24/2017 | EPI_ISL_304442 | A/Bayern/17/2018 |
| EPI_ISL_339519 | A/Saarland/28/2017 | EPI_ISL_304443 | A/Berlin/23/2018 |
| EPI_ISL_339520 | A/Saarland/29/2017 | EPI_ISL_306515 | A/Baden-Wuerttemberg/82/2018 |
| EPI_ISL_292126 | A/Saarland/26/2017 | EPI_ISL_306516 | A/Saarland/9/2018 |
| EPI_ISL_339521 | A/Saarland/30/2017 | EPI_ISL_306517 | A/Baden-Wuerttemberg/120/2018 |
| EPI_ISL_339522 | A/Saarland/31/2017 | EPI_ISL_306678 | A/Nordrhein-Westfalen/68/2018 |
| EPI_ISL_293953 | A/Nordrhein-Westfalen/138/2017 | EPI_ISL_306518 | A/Berlin/37/2018 |
| EPI_ISL_293954 | A/Baden-Wuerttemberg/257/2017 | EPI_ISL_306519 | A/Baden-Wuerttemberg/153/2018 |
| EPI_ISL_293955 | A/Baden-Wuerttemberg/259/2017 | EPI_ISL_306520 | A/Baden-Wuerttemberg/154/2018 |
| EPI_ISL_293956 | A/Bayern/1/2018 | EPI_ISL_306521 | A/Nordrhein-Westfalen/65/2018 |
| EPI_ISL_293967 | A/Nordrhein-Westfalen/2/2018 | EPI_ISL_306522 | A/Berlin/55/2018 |
| EPI_ISL_293969 | A/Hessen/1/2018 | EPI_ISL_311288 | A/Baden-Wuerttemberg/156/2018 |
| EPI_ISL_293970 | A/Baden-Wuerttemberg/4/2018 | EPI_ISL_311289 | A/Brandenburg/25/2018 |
| EPI_ISL_293972 | A/Bayern/3/2018 | EPI_ISL_311306 | A/Berlin/56/2018 |
| EPI_ISL_298744 | A/Niedersachsen/1/2018 | EPI_ISL_311290 | A/Niedersachsen/115/2018 |
| EPI_ISL_298747 | A/Rheinland-Pfalz/4/2018 | EPI_ISL_403461 | A/Schleswig-Holstein/5/2019 |
| EPI_ISL_298749 | A/Berlin/5/2018 | EPI_ISL_378172 | A/Sachsen/89/2018 |
| EPI_ISL_301673 | A/Baden-Wuerttemberg/86/2018 | EPI_ISL_378171 | A/Sachsen/88/2018 |
| EPI_ISL_301674 | A/Baden-Wuerttemberg/87/2018 | EPI_ISL_378162 | A/Baden-Wuerttemberg/309/2019 |
| EPI_ISL_301675 | A/Baden-Wuerttemberg/88/2018 | EPI_ISL_378161 | A/Schleswig-Holstein/4/2019 |
| EPI_ISL_301676 | A/Baden-Wuerttemberg/89/2018 | EPI_ISL_378160 | A/Bayern/112/2019 |
| EPI_ISL_301677 | A/Baden-Wuerttemberg/90/2018 | EPI_ISL_378159 | A/Baden-Wuerttemberg/308/2019 |
| EPI_ISL_301678 | A/Baden-Wuerttemberg/24/2018 | EPI_ISL_378158 | A/Baden-Wuerttemberg/53/2019 |
| EPI_ISL_301679 | A/Baden-Wuerttemberg/15/2018 | EPI_ISL_378157 | A/Sachsen/28/2019 |
| EPI_ISL_356688 | A/Sachsen/130/2019 | EPI_ISL_378156 | A/Niedersachsen/194/2019 |
| EPI_ISL_378155 | A/Baden-Wuerttemberg/25/2019 | EPI_ISL_346525 | A/Hessen/40/2019 |
| EPI_ISL_378154 | A/Niedersachsen/28/2019 | EPI_ISL_346524 | A/Niedersachsen/120/2019 |
| EPI_ISL_378153 | A/Berlin/7/2019 | EPI_ISL_346523 | A/Niedersachsen/66/2019 |
| EPI_ISL_356689 | A/Thueringen/107/2019 | EPI_ISL_346522 | A/Bayern/88/2019 |
| EPI_ISL_346521 | A/Saarland/6/2019 | EPI_ISL_346520 | A/Bayern/63/2019 |

Table 1S. Accession number (Gisaid) of influenza A(H1N1)pdm09 viruses isolated in Germany between 2009 and 2024

| | | | |
|---|---|---|---|
| EPI_ISL_356687 | A/Nordrhein-Westfalen/120/2019 | EPI_ISL_339515 | A/Nordrhein-Westfalen/9/2019 |
| EPI_ISL_356686 | A/Bayern/109/2019 | EPI_ISL_346519 | A/Nordrhein-Westfalen/56/2019 |
| EPI_ISL_355970 | A/Hamburg/4/2019 | EPI_ISL_346518 | A/Niedersachsen/119/2019 |
| EPI_ISL_355968 | A/Sachsen-Anhalt/84/2019 | EPI_ISL_346517 | A/Baden-Wuerttemberg/55/2019 |
| EPI_ISL_355946 | A/Hessen/67/2019 | EPI_ISL_346516 | A/Niedersachsen/40/2019 |
| EPI_ISL_355945 | A/Hessen/64/2019 | EPI_ISL_343574 | A/Berlin/26/2019 |
| EPI_ISL_355944 | A/Bayern/107/2019 | EPI_ISL_343573 | A/Thueringen/46/2019 |
| EPI_ISL_355943 | A/Sachsen/111/2019 | EPI_ISL_343572 | A/Bayern/32/2019 |
| EPI_ISL_355942 | A/Niedersachsen/160/2019 | EPI_ISL_343571 | A/Thueringen/11/2019 |
| EPI_ISL_355941 | A/Rheinland-Pfalz/51/2019 | EPI_ISL_343570 | A/Hessen/17/2019 |
| EPI_ISL_353318 | A/Baden-Wuerttemberg/195/2019 | EPI_ISL_343569 | A/Niedersachsen/26/2019 |
| EPI_ISL_353317 | A/Niedersachsen/150/2019 | EPI_ISL_343568 | A/Rheinland-Pfalz/31/2019 |
| EPI_ISL_353316 | A/Bremen/19/2019 | EPI_ISL_343567 | A/Hessen/16/2019 |
| EPI_ISL_351421 | A/Baden-Wuerttemberg/164/2019 | EPI_ISL_343566 | A/Bremen/7/2019 |
| EPI_ISL_351420 | A/Sachsen-Anhalt/43/2019 | EPI_ISL_343565 | A/Baden-Wuerttemberg/88/2019 |
| EPI_ISL_351419 | A/Brandenburg/23/2019 | EPI_ISL_343564 | A/Baden-Wuerttemberg/21/2019 |
| EPI_ISL_351418 | A/Sachsen/125/2019 | EPI_ISL_343563 | A/Berlin/45/2019 |
| EPI_ISL_351417 | A/Sachsen/124/2019 | EPI_ISL_343562 | A/Baden-Wuerttemberg/15/2019 |
| EPI_ISL_348805 | A/Baden-Wuerttemberg/212/2019 | EPI_ISL_343561 | A/Saarland/3/2019 |
| EPI_ISL_348793 | A/Sachsen/81/2019 | EPI_ISL_343560 | A/Nordrhein-Westfalen/16/2019 |
| EPI_ISL_348792 | A/Sachsen-Anhalt/40/2019 | EPI_ISL_343559 | A/Berlin/11/2019 |
| EPI_ISL_348791 | A/Niedersachsen/98/2019 | EPI_ISL_343558 | A/Rheinland-Pfalz/6/2019 |
| EPI_ISL_348790 | A/Bayern/102/2019 | EPI_ISL_343557 | A/Berlin/10/2019 |
| EPI_ISL_348789 | A/Niedersachsen/86/2019 | EPI_ISL_343556 | A/Nordrhein-Westfalen/12/2019 |
| EPI_ISL_348788 | A/Sachsen/109/2019 | EPI_ISL_339517 | A/Hessen/23/2019 |
| EPI_ISL_348786 | A/Saarland/8/2019 | EPI_ISL_339516 | A/Niedersachsen/11/2019 |
| EPI_ISL_348785 | A/Bayern/101/2019 | EPI_ISL_482809 | A/Nordrhein-Westfalen/37/2020 |
| EPI_ISL_339513 | A/Berlin/6/2019 | EPI_ISL_645137 | A/Berlin/31/2020 |
| EPI_ISL_339512 | A/Brandenburg/2/2019 | EPI_ISL_491173 | A/Hessen/47/2020 |
| EPI_ISL_339510 | A/Sachsen-Anhalt/1/2019 | EPI_ISL_482847 | A/Brandenburg/8/2020 |
| EPI_ISL_339509 | A/Niedersachsen/30/2019 | EPI_ISL_482810 | A/Nordrhein-Westfalen/59/2020 |
| EPI_ISL_339508 | A/Brandenburg/1/2019 | EPI_ISL_482808 | A/Rheinland-Pfalz/18/2020 |

Table 1S. Accession number (Gisaid) of influenza A(H1N1)pdm09 viruses isolated in Germany between 2009 and 2024

| | | | |
|---|---|---|---|
| EPI_ISL_339506 | A/Nordrhein-Westfalen/5/2019 | EPI_ISL_481389 | A/Hessen/23/2020 |
| EPI_ISL_339505 | A/Rheinland-Pfalz/41/2018 | EPI_ISL_482807 | A/Hessen/30/2020 |
| EPI_ISL_339504 | A/Rheinland-Pfalz/16/2019 | EPI_ISL_482806 | A/Baden-Wuerttemberg/121/2020 |
| EPI_ISL_337000 | A/Rheinland-Pfalz/2/2019 | EPI_ISL_482805 | A/Niedersachsen/52/2020 |
| EPI_ISL_336996 | A/Baden-Wuerttemberg/4/2019 | EPI_ISL_482804 | A/Baden-Wuerttemberg/120/2020 |
| EPI_ISL_336995 | A/Bayern/2/2019 | EPI_ISL_482803 | A/Rheinland-Pfalz/24/2020 |
| EPI_ISL_336994 | A/Rheinland-Pfalz/1/2019 | EPI_ISL_482802 | A/Thueringen/17/2020 |
| EPI_ISL_336993 | A/Thueringen/1/2019 | EPI_ISL_482801 | A/Bayern/43/2020 |
| EPI_ISL_336992 | A/Nordrhein-Westfalen/1/2019 | EPI_ISL_482800 | A/Bayern/26/2020 |
| EPI_ISL_336991 | A/Bremen/1/2019 | EPI_ISL_482799 | A/Sachsen/28/2020 |
| EPI_ISL_336987 | A/ Nordrhein-Westfalen/8/2019 | EPI_ISL_482798 | A/Mecklenburg-Vorpommern/3/2020 |
| EPI_ISL_336985 | A/Niedersachsen/1/2019 | EPI_ISL_482797 | A/Bayern/25/2020 |
| EPI_ISL_336984 | A/Bayern/1/2019 | EPI_ISL_482796 | A/Schleswig-Holstein/4/2020 |
| EPI_ISL_335316 | A/Mecklenburg-Vorpommern/5/2018 | EPI_ISL_482795 | A/Baden-Wuerttemberg/119/2020 |
| EPI_ISL_335310 | A/Hamburg/7/2018 | EPI_ISL_482794 | A/Niedersachsen/49/2020 |
| EPI_ISL_335308 | A/Nordrhein-Westfalen/80/2018 | EPI_ISL_482793 | A/Baden-Wuerttemberg/58/2020 |
| EPI_ISL_335307 | A/Hamburg/6/2018 | EPI_ISL_482792 | A/Bremen/4/2020 |
| EPI_ISL_335306 | A/Berlin/58/2018 | EPI_ISL_482791 | A/Thueringen/13/2020 |
| EPI_ISL_335304 | A/Bayern/51/2018 | EPI_ISL_482790 | A/Thueringen/14/2020 |
| EPI_ISL_335303 | A/Bayern/50/2018 | EPI_ISL_482789 | A/Saarland/4/2020 |
| EPI_ISL_332946 | A/Mecklenburg-Vorpommern/4/2018 | EPI_ISL_482788 | A/Berlin/30/2020 |
| EPI_ISL_332944 | A/Baden-Wuerttemberg/163/2018 | EPI_ISL_482787 | A/Bayern/42/2020 |
| EPI_ISL_332942 | A/Thueringen/61/2018 | EPI_ISL_482786 | A/Brandenburg/10/2020 |
| EPI_ISL_332941 | A/Baden-Wuerttemberg/162/2018 | EPI_ISL_481430 | A/Bremen/13/2020 |
| EPI_ISL_645140 | A/Rheinland-Pfalz/35/2020 | EPI_ISL_481429 | A/Hessen/39/2020 |
| EPI_ISL_645139 | A/Berlin/47/2020 | EPI_ISL_481428 | A/Baden-Wuerttemberg/149/2020 |
| EPI_ISL_645138 | A/Hamburg/1/2020 | EPI_ISL_481427 | A/Nordrhein-Westfalen/77/2020 |
| EPI_ISL_481426 | A/Niedersachsen/137/2020 | EPI_ISL_481394 | A/Bayern/37/2020 |
| EPI_ISL_481425 | A/Niedersachsen/98/2020 | EPI_ISL_481393 | A/Nordrhein-Westfalen/45/2020 |
| EPI_ISL_481424 | A/Bayern/49/2020 | EPI_ISL_481391 | A/Mecklenburg-Vorpommern/4/2020 |
| EPI_ISL_481423 | A/Rheinland-Pfalz/32/2020 | EPI_ISL_481390 | A/Baden-Wuerttemberg/175/2020 |
| EPI_ISL_481422 | A/Saarland/9/2020 | EPI_ISL_481388 | A/Bayern/57/2020 |

Table 1S. Accession number (Gisaid) of influenza A(H1N1)pdm09 viruses isolated in Germany between 2009 and 2024

| | | | |
|---|---|---|---|
| EPI_ISL_481421 | A/Nordrhein-Westfalen/76/2020 | EPI_ISL_407178 | A/Bremen/1/2020 |
| EPI_ISL_481420 | A/Nordrhein-Westfalen/61/2020 | EPI_ISL_481387 | A/Saarland/6/2020 |
| EPI_ISL_481419 | A/Bayern/46/2020 | EPI_ISL_481386 | A/Hessen/22/2020 |
| EPI_ISL_481418 | A/Rheinland-Pfalz/31/2020 | EPI_ISL_481385 | A/Berlin/67/2020 |
| EPI_ISL_481417 | A/Bremen/11/2020 | EPI_ISL_481384 | A/Berlin/39/2020 |
| EPI_ISL_481416 | A/Rheinland-Pfalz/29/2020 | EPI_ISL_481383 | A/Saarland/5/2020 |
| EPI_ISL_481415 | A/Brandenburg/11/2020 | EPI_ISL_481382 | A/Baden-Wuerttemberg/85/2020 |
| EPI_ISL_481414 | A/Baden-Wuerttemberg/139/2020 | EPI_ISL_481381 | A/Nordrhein-Westfalen/78/2020 |
| EPI_ISL_481413 | A/Hessen/43/2020 | EPI_ISL_413430 | A/Baden-Wuerttemberg/49/2020 |
| EPI_ISL_481412 | A/Hessen/42/2020 | EPI_ISL_413429 | A/Nordrhein-Westfalen/47/2020 |
| EPI_ISL_481411 | A/Niedersachsen/91/2020 | EPI_ISL_413428 | A/Nordrhein-Westfalen/27/2020 |
| EPI_ISL_481410 | A/Bayern/54/2020 | EPI_ISL_413427 | A/Baden-Wuerttemberg/109/2020 |
| EPI_ISL_481409 | A/Bayern/55/2020 | EPI_ISL_413426 | A/Baden-Wuerttemberg/47/2020 |
| EPI_ISL_481408 | A/Bayern/53/2020 | EPI_ISL_413425 | A/Niedersachsen/21/2020 |
| EPI_ISL_481407 | A/Baden-Wuerttemberg/176/2020 | EPI_ISL_413424 | A/Saarland/3/2020 |
| EPI_ISL_481406 | A/Niedersachsen/87/2020 | EPI_ISL_413423 | A/Rheinland-Pfalz/12/2020 |
| EPI_ISL_481405 | A/Saarland/7/2020 | EPI_ISL_413422 | A/Sachsen/14/2020 |
| EPI_ISL_481404 | A/Rheinland-Pfalz/21/2020 | EPI_ISL_413421 | A/Berlin/24/2020 |
| EPI_ISL_481403 | A/Berlin/43/2020 | EPI_ISL_413420 | A/Baden-Wuerttemberg/28/2020 |
| EPI_ISL_481402 | A/Schleswig-Holstein/7/2020 | EPI_ISL_413419 | A/Bayern/19/2020 |
| EPI_ISL_481401 | A/Rheinland-Pfalz/36/2020 | EPI_ISL_410964 | A/Nordrhein-Westfalen/19/2020 |
| EPI_ISL_481400 | A/Nordrhein-Westfalen/79/2020 | EPI_ISL_410963 | A/Baden-Wuerttemberg/22/2020 |
| EPI_ISL_481399 | A/Nordrhein-Westfalen/44/2020 | EPI_ISL_410962 | A/Hessen/18/2020 |
| EPI_ISL_481398 | A/Berlin/68/2020 | EPI_ISL_410961 | A/Niedersachsen/16/2020 |
| EPI_ISL_481397 | A/Nordrhein-Westfalen/46/2020 | EPI_ISL_410960 | A/Berlin/15/2020 |
| EPI_ISL_481396 | A/Hessen/41/2020 | EPI_ISL_410959 | A/Baden-Wuerttemberg/21/2020 |
| EPI_ISL_481395 | A/Hessen/24/2020 | EPI_ISL_410958 | A/Bayern/16/2020 |
| EPI_ISL_410957 | A/Bayern/15/2020 | EPI_ISL_407182 | A/Mecklenburg-Vorpommern/1/2020 |
| EPI_ISL_410956 | A/Sachsen/10/2020 | EPI_ISL_407181 | A/Baden-Wuerttemberg/1/2020 |
| EPI_ISL_410955 | A/Bayern/14/2020 | EPI_ISL_407180 | A/Rheinland-Pfalz/3/2020 |
| EPI_ISL_410954 | A/Baden-Wuerttemberg/19/2020 | EPI_ISL_407179 | A/Brandenburg/2/2020 |
| EPI_ISL_410953 | A/Baden-Wuerttemberg/18/2020 | EPI_ISL_407177 | A/Saarland/1/2020 |

Table 1S. Accession number (Gisaid) of influenza A(H1N1)pdm09 viruses isolated in Germany between 2009 and 2024

| | | | |
|---|---|---|---|
| EPI_ISL_410952 | A/Bayern/9/2020 | EPI_ISL_401777 | A/Hessen/71/2019 |
| EPI_ISL_410951 | A/Nordrhein-Westfalen/14/2020 | EPI_ISL_407176 | A/Brandenburg/1/2020 |
| EPI_ISL_410950 | A/Baden-Wuerttemberg/11/2020 | EPI_ISL_407175 | A/Nordrhein-Westfalen/2/2020 |
| EPI_ISL_410949 | A/Hessen/5/2020 | EPI_ISL_407174 | A/Thueringen/1/2020 |
| EPI_ISL_410075 | A/Niedersachsen/11/2020 | EPI_ISL_407173 | A/Nordrhein-Westfalen/1/2020 |
| EPI_ISL_410073 | A/Baden-Wuerttemberg/5/2020 | EPI_ISL_407172 | A/Rheinland-Pfalz/61/2019 |
| EPI_ISL_410071 | A/Mecklenburg-Vorpommern/2/2020 | EPI_ISL_407171 | A/Rheinland-Pfalz/60/2019 |
| EPI_ISL_410070 | A/Hessen/3/2020 | EPI_ISL_407170 | A/Schleswig-Holstein/6/2019 |
| EPI_ISL_410068 | A/Nordrhein-Westfalen/11/2020 | EPI_ISL_407169 | A/Schleswig-Holstein/7/2019 |
| EPI_ISL_410066 | A/Hessen/2/2020 | EPI_ISL_405006 | A/Nordrhein-Westfalen/143/2019 |
| EPI_ISL_410065 | A/Berlin/12/2020 | EPI_ISL_405005 | A/Thueringen/125/2019 |
| EPI_ISL_410063 | A/Nordrhein-Westfalen/9/2020 | EPI_ISL_405004 | A/Nordrhein-Westfalen/142/2019 |
| EPI_ISL_410062 | A/Niedersachsen/9/2020 | EPI_ISL_405003 | A/Mecklenburg-Vorpommern/13/2019 |
| EPI_ISL_410060 | A/Nordrhein-Westfalen/8/2020 | EPI_ISL_405002 | A/Hessen/73/2019 |
| EPI_ISL_410059 | A/Nordrhein-Westfalen/7/2020 | EPI_ISL_405001 | A/Mecklenburg-Vorpommern/12/2019 |
| EPI_ISL_410058 | A/Nordrhein-Westfalen/6/2020 | EPI_ISL_405000 | A/Nordrhein-Westfalen/139/2019 |
| EPI_ISL_410057 | A/Bayern/7/2020 | EPI_ISL_404999 | A/Brandenburg/29/2019 |
| EPI_ISL_410056 | A/Bayern/5/2020 | EPI_ISL_404998 | A/Nordrhein-Westfalen/138/2019 |
| EPI_ISL_410055 | A/Niedersachsen/4/2020 | EPI_ISL_404997 | A/Baden-Wuerttemberg/313/2019 |
| EPI_ISL_410054 | A/Niedersachsen/2/2020 | EPI_ISL_404996 | A/Sachsen/152/2019 |
| EPI_ISL_410053 | A/Berlin/6/2020 | EPI_ISL_404995 | A/Sachsen/151/2019 |
| EPI_ISL_407188 | A/Sachsen/3/2020 | EPI_ISL_404994 | A/Nordrhein-Westfalen/136/2019 |
| EPI_ISL_407187 | A/Nordrhein-Westfalen/4/2020 | EPI_ISL_401781 | A/Berlin/59/2019 |
| EPI_ISL_407186 | A/Hessen/7/2020 | EPI_ISL_401780 | A/Berlin/58/2019 |
| EPI_ISL_407185 | A/Bayern/1/2020 | EPI_ISL_401779 | A/Niedersachsen/195/2019 |
| EPI_ISL_407184 | A/Thueringen/4/2020 | EPI_ISL_401778 | A/Berlin/57/2019 |
| EPI_ISL_407183 | A/Nordrhein-Westfalen/3/2020 | EPI_ISL_17780387 | A/Niedersachsen/9/2023 |
| EPI_ISL_2434153 | A/Mecklenburg-Vorpommern/1-A/2021 | EPI_ISL_17959494 | A/Baden-Wuerttemberg/27/2023 |
| EPI_ISL_2434152 | A/Mecklenburg-Vorpommern/1/2021 | EPI_ISL_17959493 | A/Hessen/11/2023 |
| EPI_ISL_10429833 | A/Mecklenburg-Vorpommern/2/2022 | EPI_ISL_17959492 | A/Berlin/19/2023 |
| EPI_ISL_11789577 | A/Bremen/3/2022 | EPI_ISL_17959491 | A/Rheinland-Pfalz/6/2023 |
| EPI_ISL_13503236 | A/Rheinland-Pfalz/2/2022 | EPI_ISL_17780386 | A/Brandenburg/2/2023 |

Table 1S. Accession number (Gisaid) of influenza A(H1N1)pdm09 viruses isolated in Germany between 2009 and 2024

| | | | |
|---|---|---|---|
| EPI_ISL_12839598 | A/Bremen/10/2022 | EPI_ISL_17592912 | A/Bayern/4/2023 |
| EPI_ISL_12839599 | A/Hessen/16/2022 | EPI_ISL_17780385 | A/Hessen/7/2023 |
| EPI_ISL_13503237 | A/Saarland/2/2022 | EPI_ISL_17780384 | A/Thueringen/5/2023 |
| EPI_ISL_13638360 | A/Niedersachsen/12/2022 | EPI_ISL_17780383 | A/Baden-Wuerttemberg/13/2023 |
| EPI_ISL_14925890 | A/Baden-Wuerttemberg/30/2022 | EPI_ISL_17780382 | A/Niedersachsen/4/2023 |
| EPI_ISL_14934966 | A/Nordrhein-Westfalen/31/2022 | EPI_ISL_17780308 | A/Baden-Wuerttemberg/24/2023 |
| EPI_ISL_227814 | A/Niedersachsen/44/2022 | EPI_ISL_17780307 | A/Berlin/16/2023 |
| EPI_ISL_227814 | A/Niedersachsen/45/2022 | EPI_ISL_17780306 | A/Berlin/8/2023 |
| EPI_ISL_227814 | A/Brandenburg/6/2022 | EPI_ISL_17780305 | A/Nordrhein-Westfalen/9/2023 |
| EPI_ISL_19168065 | A/Berlin/270/2022 | EPI_ISL_17780304 | A/Baden-Wuerttemberg/20/2023 |
| EPI_ISL_19168066 | A/Berlin/271/2022 | EPI_ISL_17780303 | A/Hessen/8/2023 |
| EPI_ISL_19168067 | A/Berlin/139/2022 | EPI_ISL_17780302 | A/Hessen/4/2023 |
| EPI_ISL_19168068 | A/Hessen/37/2022 | EPI_ISL_17780301 | A/Nordrhein-Westfalen/3/2023 |
| EPI_ISL_19168069 | A/Rheinland-Pfalz/15/2022 | EPI_ISL_17592920 | A/Berlin/12/2023 |
| EPI_ISL_18526252 | A/Berlin/22/2023 | EPI_ISL_17592919 | A/Berlin/6/2023 |
| EPI_ISL_18526251 | A/Niedersachsen/21/2023 | EPI_ISL_17592918 | A/Nordrhein-Westfalen/8/2023 |
| EPI_ISL_18526250 | A/Berlin/14/2023 | EPI_ISL_17592917 | A/Sachsen/3/2023 |
| EPI_ISL_18514884 | A/Berlin/26/2023 | EPI_ISL_17592916 | A/Berlin/2/2023 |
| EPI_ISL_18412346 | A/Nordrhein-Westfalen/10/2023 | EPI_ISL_17592915 | A/Bayern/7/2023 |
| EPI_ISL_18412345 | A/Sachsen-Anhalt/8/2023 | EPI_ISL_17592914 | A/Saarland/1/2023 |
| EPI_ISL_18412344 | A/Baden-Wuerttemberg/27/2023 | EPI_ISL_17592913 | A/Berlin/11/2023 |
| EPI_ISL_17959500 | A/Schleswig-Holstein/2/2023 | EPI_ISL_17197521 | A/Schweiz/1/2023 |
| EPI_ISL_17959499 | A/Berlin/21/2023 | EPI_ISL_16548602 | A/Berlin/220/2022 |
| EPI_ISL_17959498 | A/Rheinland-Pfalz/5/2023 | EPI_ISL_17197520 | A/Baden-Wuerttemberg/158/2022 |
| EPI_ISL_17959497 | A/Hessen/13/2023 | EPI_ISL_17037583 | A/Nordrhein-Westfalen/2/2023 |
| EPI_ISL_17959496 | A/Hessen/12/2023 | EPI_ISL_17037582 | A/Nordrhein-Westfalen/6/2023 |
| EPI_ISL_17959495 | A/Berlin/20/2023 | EPI_ISL_17037581 | A/Nordrhein-Westfalen/1/2023 |
| EPI_ISL_17037580 | A/Mecklenburg-Vorpommern/1/2023 | EPI_ISL_17037548 | A/Nordrhein-Westfalen/58/2022 |
| EPI_ISL_17037579 | A/Bayern/1/2023 | EPI_ISL_16548605 | A/Hamburg/4/2022 |
| EPI_ISL_17037578 | A/Thueringen/4/2023 | EPI_ISL_16548604 | A/Baden-Wuerttemberg/108/2022 |
| EPI_ISL_17037577 | A/Baden-Wuerttemberg/1/2023 | EPI_ISL_16548603 | A/Nordrhein-Westfalen/90/2022 |
| EPI_ISL_17037576 | A/Bayern/48/2022 | EPI_ISL_16548601 | A/Thueringen/46/2022 |

Table 1S. Accession number (Gisaid) of influenza A(H1N1)pdm09 viruses isolated in Germany between 2009 and 2024

| | | | |
|---|---|---|---|
| EPI_ISL_17037575 | A/Berlin/228/2022 | EPI_ISL_16103551 | A/Nordrhein-Westfalen/39/2022 |
| EPI_ISL_17037574 | A/Baden-Wuerttemberg/96/2022 | EPI_ISL_16548599 | A/Thueringen/81/2022 |
| EPI_ISL_17037573 | A/Baden-Wuerttemberg/95/2022 | EPI_ISL_16103552 | A/Schleswig-Holstein/19/2022 |
| EPI_ISL_17037572 | A/Berlin/211/2022 | EPI_ISL_18993828 | A/Nordrhein-Westfalen/67/2024 |
| EPI_ISL_17037571 | A/Nordrhein-Westfalen/88/2022 | EPI_ISL_19002011 | A/Schleswig-Holstein/44/2024 |
| EPI_ISL_17037570 | A/Berlin/209/2022 | EPI_ISL_19002010 | A/Schleswig-Holstein/43/2024 |
| EPI_ISL_17037569 | A/Nordrhein-Westfalen/85/2022 | EPI_ISL_19002009 | A/Schleswig-Holstein/42/2024 |
| EPI_ISL_17037568 | A/Nordrhein-Westfalen/87/2022 | EPI_ISL_19002008 | A/Schleswig-Holstein/41/2024 |
| EPI_ISL_17037567 | A/Berlin/227/2022 | EPI_ISL_19002007 | A/Schleswig-Holstein/28/2024 |
| EPI_ISL_17037566 | A/Sachsen/30/2022 | EPI_ISL_19002006 | A/Schleswig-Holstein/40/2024 |
| EPI_ISL_17037565 | A/Baden-Wuerttemberg/94/2022 | EPI_ISL_19002005 | A/Schleswig-Holstein/39/2024 |
| EPI_ISL_17037564 | A/Rheinland-Pfalz/36/2022 | EPI_ISL_19002004 | A/Schleswig-Holstein/24/2024 |
| EPI_ISL_17037563 | A/Thueringen/73/2022 | EPI_ISL_19002003 | A/Thueringen/63/2024 |
| EPI_ISL_17037562 | A/Baden-Wuerttemberg/92/2022 | EPI_ISL_19002002 | A/Nordrhein-Westfalen/72/2024 |
| EPI_ISL_17037561 | A/Niedersachsen/106/2022 | EPI_ISL_19002001 | A/Nordrhein-Westfalen/71/2024 |
| EPI_ISL_17037560 | A/Berlin/203/2022 | EPI_ISL_19002000 | A/Hessen/80/2024 |
| EPI_ISL_17037559 | A/Berlin/202/2022 | EPI_ISL_19001999 | A/Hessen/79/2024 |
| EPI_ISL_17037558 | A/Berlin/201/2022 | EPI_ISL_19001998 | A/Bremen/21/2024 |
| EPI_ISL_17037557 | A/Berlin/198/2022 | EPI_ISL_19001997 | A/Bremen/20/2024 |
| EPI_ISL_17037556 | A/Berlin/196/2022 | EPI_ISL_19001996 | A/Hessen/78/2024 |
| EPI_ISL_17037555 | A/Berlin/230/2022 | EPI_ISL_19001995 | A/Rheinland-Pfalz/31/2024 |
| EPI_ISL_17037554 | A/Nordrhein-Westfalen/76/2022 | EPI_ISL_19001994 | A/Baden-Wuerttemberg/136/2024 |
| EPI_ISL_17037553 | A/Berlin/181/2022 | EPI_ISL_19001993 | A/Hessen/77/2024 |
| EPI_ISL_17037552 | A/Berlin/178/2022 | EPI_ISL_19001992 | A/Nordrhein-Westfalen/70/2024 |
| EPI_ISL_17037551 | A/Nordrhein-Westfalen/59/2022 | EPI_ISL_19001991 | A/Nordrhein-Westfalen/69/2024 |
| EPI_ISL_17037550 | A/Berlin/226/2022 | EPI_ISL_19001990 | A/Saarland/15/2024 |
| EPI_ISL_17037549 | A/Berlin/224/2022 | EPI_ISL_19001989 | A/Nordrhein-Westfalen/68/2024 |
| EPI_ISL_19001988 | A/Schleswig-Holstein/23/2024 | EPI_ISL_18993832 | A/Bayern/61/2024 |
| EPI_ISL_19001987 | A/Hessen/76/2024 | EPI_ISL_18993831 | A/Hessen/69/2024 |
| EPI_ISL_19001986 | A/Baden-Wuerttemberg/135/2024 | EPI_ISL_18993830 | A/Hessen/68/2024 |
| EPI_ISL_19001985 | A/Niedersachsen/75/2024 | EPI_ISL_18993829 | A/Hessen/67/2024 |
| EPI_ISL_19001984 | A/Berlin/30/2024 | EPI_ISL_18993827 | A/Bayern/60/2024 |

Table 1S. Accession number (Gisaid) of influenza A(H1N1)pdm09 viruses isolated in Germany between 2009 and 2024

| | | | |
|---|---|---|---|
| EPI_ISL_19001983 | A/Bayern/69/2024 | EPI_ISL_18969203 | A/Hessen/59/2024 |
| EPI_ISL_19001982 | A/Bayern/68/2024 | EPI_ISL_18993826 | A/Bayern/59/2024 |
| EPI_ISL_19001981 | A/Bayern/67/2024 | EPI_ISL_18993825 | A/Bremen/19/2024 |
| EPI_ISL_19001980 | A/Bayern/66/2024 | EPI_ISL_18993824 | A/Schleswig-Holstein/37/2024 |
| EPI_ISL_19001979 | A/Hessen/75/2024 | EPI_ISL_18993823 | A/Niedersachsen/96/2024 |
| EPI_ISL_19001978 | A/Hessen/74/2024 | EPI_ISL_18993822 | A/Baden-Wuerttemberg/131/2024 |
| EPI_ISL_19001977 | A/Bayern/65/2024 | EPI_ISL_18993821 | A/Hessen/66/2024 |
| EPI_ISL_19001976 | A/Bayern/64/2024 | EPI_ISL_18993820 | A/Bayern/58/2024 |
| EPI_ISL_19001975 | A/Bayern/63/2024 | EPI_ISL_18993819 | A/Bayern/57/2024 |
| EPI_ISL_19001974 | A/Mecklenburg-Vorpommern/15/2024 | EPI_ISL_18993818 | A/Bayern/56/2024 |
| EPI_ISL_19001973 | A/Mecklenburg-Vorpommern/11/2024 | EPI_ISL_18993817 | A/Hessen/65/2024 |
| EPI_ISL_18993848 | A/Schleswig-Holstein/38/2024 | EPI_ISL_18993816 | A/Hessen/64/2024 |
| EPI_ISL_18993847 | A/Brandenburg/27/2024 | EPI_ISL_18993815 | A/Brandenburg/17/2024 |
| EPI_ISL_18993846 | A/Niedersachsen/98/2024 | EPI_ISL_18993814 | A/Brandenburg/26/2024 |
| EPI_ISL_18993845 | A/Niedersachsen/97/2024 | EPI_ISL_18993813 | A/Thueringen/61/2024 |
| EPI_ISL_18993844 | A/Bayern/62/2024 | EPI_ISL_18993812 | A/Baden-Wuerttemberg/130/2024 |
| EPI_ISL_18993843 | A/Baden-Wuerttemberg/134/2024 | EPI_ISL_18993811 | A/Bremen/15/2024 |
| EPI_ISL_18993842 | A/Baden-Wuerttemberg/133/2024 | EPI_ISL_18993810 | A/Hessen/63/2024 |
| EPI_ISL_18993841 | A/Rheinland-Pfalz/30/2024 | EPI_ISL_18993809 | A/Rheinland-Pfalz/28/2024 |
| EPI_ISL_18993840 | A/Saarland/14/2024 | EPI_ISL_18993808 | A/Niedersachsen/49/2024 |
| EPI_ISL_18993839 | A/Hessen/73/2024 | EPI_ISL_18993807 | A/Niedersachsen/95/2024 |
| EPI_ISL_18993838 | A/Hessen/72/2024 | EPI_ISL_18993806 | A/Mecklenburg-Vorpommern/14/2024 |
| EPI_ISL_18993837 | A/Rheinland-Pfalz/29/2024 | EPI_ISL_18983414 | A/Bremen/18/2024 |
| EPI_ISL_18993836 | A/Baden-Wuerttemberg/132/2024 | EPI_ISL_18983413 | A/Sachsen-Anhalt/26/2024 |
| EPI_ISL_18993835 | A/Thueringen/62/2024 | EPI_ISL_18983412 | A/Thueringen/57/2024 |
| EPI_ISL_18993834 | A/Hessen/71/2024 | EPI_ISL_18983411 | A/Thueringen/56/2024 |
| EPI_ISL_18993833 | A/Hessen/70/2024 | EPI_ISL_18983410 | A/Thueringen/55/2024 |
| EPI_ISL_18983409 | A/Rheinland-Pfalz/16/2024 | EPI_ISL_18976589 | A/Niedersachsen/79/2024 |
| EPI_ISL_18983408 | A/Rheinland-Pfalz/15/2024 | EPI_ISL_18976588 | A/Nordrhein-Westfalen/58/2024 |
| EPI_ISL_18983407 | A/Thueringen/54/2024 | EPI_ISL_18976587 | A/Bayern/48/2024 |
| EPI_ISL_18983406 | A/Baden-Wuerttemberg/122/2024 | EPI_ISL_18969204 | A/Nordrhein-Westfalen/55/2024 |
| EPI_ISL_18983405 | A/Hessen/62/2024 | EPI_ISL_18969202 | A/Hessen/58/2024 |

Table 1S. Accession number (Gisaid) of influenza A(H1N1)pdm09 viruses isolated in Germany between 2009 and 2024

| | | | |
|---|---|---|---|
| EPI_ISL_18983404 | A/Bayern/54/2024 | EPI_ISL_18952709 | A/Nordrhein-Westfalen/49/2024 |
| EPI_ISL_18983403 | A/Mecklenburg-Vorpommern/10/2024 | EPI_ISL_18969201 | A/Rheinland-Pfalz/27/2024 |
| EPI_ISL_18983402 | A/Baden-Wuerttemberg/65/2024 | EPI_ISL_18969200 | A/Rheinland-Pfalz/26/2024 |
| EPI_ISL_18983401 | A/Baden-Wuerttemberg/121/2024 | EPI_ISL_18969199 | A/Hessen/57/2024 |
| EPI_ISL_18983400 | A/Thueringen/27/2024 | EPI_ISL_18969198 | A/Baden-Wuerttemberg/111/2024 |
| EPI_ISL_18983399 | A/Bayern/53/2024 | EPI_ISL_18969197 | A/Bayern/21/2024 |
| EPI_ISL_18983398 | A/Nordrhein-Westfalen/41/2024 | EPI_ISL_18969196 | A/Niedersachsen/78/2024 |
| EPI_ISL_18983397 | A/Bayern/52/2024 | EPI_ISL_18969195 | A/Baden-Wuerttemberg/110/2024 |
| EPI_ISL_18983396 | A/Berlin/29/2024 | EPI_ISL_18969194 | A/Rheinland-Pfalz/25/2024 |
| EPI_ISL_18983395 | A/Hessen/34/2024 | EPI_ISL_18969193 | A/Hessen/56/2024 |
| EPI_ISL_18983394 | A/Hessen/46/2024 | EPI_ISL_18969192 | A/Baden-Wuerttemberg/109/2024 |
| EPI_ISL_18983393 | A/Nordrhein-Westfalen/62/2024 | EPI_ISL_18969191 | A/Baden-Wuerttemberg/108/2024 |
| EPI_ISL_18983392 | A/Nordrhein-Westfalen/39/2024 | EPI_ISL_18969190 | A/Hessen/55/2024 |
| EPI_ISL_18983391 | A/Bayern/51/2024 | EPI_ISL_18969189 | A/Hessen/54/2024 |
| EPI_ISL_18983390 | A/Bayern/39/2024 | EPI_ISL_18969188 | A/Hessen/22/2024 |
| EPI_ISL_18983389 | A/Nordrhein-Westfalen/61/2024 | EPI_ISL_18969187 | A/Hessen/53/2024 |
| EPI_ISL_18983387 | A/Schleswig-Holstein/36/2024 | EPI_ISL_18969186 | A/Nordrhein-Westfalen/54/2024 |
| EPI_ISL_18983386 | A/Schleswig-Holstein/35/2024 | EPI_ISL_18969185 | A/Nordrhein-Westfalen/53/2024 |
| EPI_ISL_18983385 | A/Schleswig-Holstein/34/2024 | EPI_ISL_18969184 | A/Mecklenburg-Vorpommern/12/2024 |
| EPI_ISL_18983384 | A/Schleswig-Holstein/33/2024 | EPI_ISL_18969183 | A/Baden-Wuerttemberg/107/2024 |
| EPI_ISL_18983383 | A/Schleswig-Holstein/32/2024 | EPI_ISL_18969182 | A/Baden-Wuerttemberg/106/2024 |
| EPI_ISL_18983382 | A/Schleswig-Holstein/31/2024 | EPI_ISL_18969181 | A/Bayern/45/2024 |
| EPI_ISL_18983381 | A/Baden-Wuerttemberg/84/2024 | EPI_ISL_18969180 | A/Bayern/44/2024 |
| EPI_ISL_18983380 | A/Baden-Wuerttemberg/112/2024 | EPI_ISL_18969179 | A/Bayern/43/2024 |
| EPI_ISL_18976592 | A/Nordrhein-Westfalen/60/2024 | EPI_ISL_18969178 | A/Baden-Wuerttemberg/105/2024 |
| EPI_ISL_18976591 | A/Nordrhein-Westfalen/59/2024 | EPI_ISL_18969177 | A/Saarland/13/2024 |
| EPI_ISL_18976590 | A/Bayern/49/2024 | EPI_ISL_18969176 | A/Baden-Wuerttemberg/104/2024 |
| EPI_ISL_18969175 | A/Baden-Wuerttemberg/103/2024 | EPI_ISL_18952714 | A/Nordrhein-Westfalen/51/2024 |
| EPI_ISL_18969174 | A/Hessen/52/2024 | EPI_ISL_18952713 | A/Thueringen/41/2024 |
| EPI_ISL_18969173 | A/Hessen/51/2024 | EPI_ISL_18952712 | A/Rheinland-Pfalz/24/2024 |
| EPI_ISL_18969172 | A/Hessen/50/2024 | EPI_ISL_18952711 | A/Rheinland-Pfalz/23/2024 |
| EPI_ISL_18969171 | A/Hessen/49/2024 | EPI_ISL_18952708 | A/Baden-Wuerttemberg/52/2024 |

Table 1S. Accession number (Gisaid) of influenza A(H1N1)pdm09 viruses isolated in Germany between 2009 and 2024

| | | | |
|---|---|---|---|
| EPI_ISL_18969170 | A/Nordrhein-Westfalen/52/2024 | EPI_ISL_18941643 | A/Thueringen/40/2024 |
| EPI_ISL_18969169 | A/Thueringen/44/2024 | EPI_ISL_18952707 | A/Berlin/25/2024 |
| EPI_ISL_18969168 | A/Bayern/42/2024 | EPI_ISL_18952706 | A/Brandenburg/23/2024 |
| EPI_ISL_18969167 | A/Bremen/12/2024 | EPI_ISL_18952705 | A/Baden-Wuerttemberg/91/2024 |
| EPI_ISL_18969166 | A/Schleswig-Holstein/29/2024 | EPI_ISL_18952704 | A/Niedersachsen/76/2024 |
| EPI_ISL_18969165 | A/Baden-Wuerttemberg/102/2024 | EPI_ISL_18952703 | A/Niedersachsen/35/2024 |
| EPI_ISL_18969164 | A/Brandenburg/25/2024 | EPI_ISL_18952702 | A/Niedersachsen/34/2024 |
| EPI_ISL_18969163 | A/Baden-Wuerttemberg/101/2024 | EPI_ISL_18952701 | A/Berlin/18/2024 |
| EPI_ISL_18969162 | A/Thueringen/43/2024 | EPI_ISL_18952700 | A/Rheinland-Pfalz/22/2024 |
| EPI_ISL_18961387 | A/Baden-Wuerttemberg/100/2024 | EPI_ISL_18952699 | A/Sachsen/9/2024 |
| EPI_ISL_18961386 | A/Niedersachsen/1/2024 | EPI_ISL_18952698 | A/Hessen/48/2024 |
| EPI_ISL_18961385 | A/Brandenburg/24/2024 | EPI_ISL_18952697 | A/Hessen/47/2024 |
| EPI_ISL_18961384 | A/Bayern/1/2024 | EPI_ISL_18952696 | A/Baden-Wuerttemberg/90/2024 |
| EPI_ISL_18961383 | A/Bayern/41/2024 | EPI_ISL_18952695 | A/Baden-Wuerttemberg/89/2024 |
| EPI_ISL_18961382 | A/Thueringen/42/2024 | EPI_ISL_18952694 | A/Baden-Wuerttemberg/30/2024 |
| EPI_ISL_18961381 | A/Bayern/18/2023 | EPI_ISL_18952693 | A/Baden-Wuerttemberg/88/2024 |
| EPI_ISL_18961380 | A/Bayern/17/2023 | EPI_ISL_18952692 | A/Hessen/16/2024 |
| EPI_ISL_18961379 | A/Niedersachsen/26/2023 | EPI_ISL_18952691 | A/Baden-Wuerttemberg/87/2024 |
| EPI_ISL_18961378 | A/Hessen/47/2023 | EPI_ISL_18952690 | A/Mecklenburg-Vorpommern/7/2024 |
| EPI_ISL_18961377 | A/Bayern/21/2023 | EPI_ISL_18952689 | A/Nordrhein-Westfalen/48/2024 |
| EPI_ISL_18961376 | A/Hessen/46/2023 | EPI_ISL_18952688 | A/Rheinland-Pfalz/21/2024 |
| EPI_ISL_18961375 | A/Hessen/45/2023 | EPI_ISL_18952687 | A/Bayern/20/2024 |
| EPI_ISL_18961374 | A/Baden-Wuerttemberg/54/2023 | EPI_ISL_18952686 | A/Schleswig-Holstein/7/2024 |
| EPI_ISL_18961373 | A/Baden-Wuerttemberg/53/2023 | EPI_ISL_18952685 | A/Nordrhein-Westfalen/21/2024 |
| EPI_ISL_18961372 | A/Baden-Wuerttemberg/34/2023 | EPI_ISL_18952684 | A/Brandenburg/22/2024 |
| EPI_ISL_18961371 | A/Hessen/34/2023 | EPI_ISL_18952683 | A/Baden-Wuerttemberg/86/2024 |
| EPI_ISL_18952715 | A/Niedersachsen/36/2024 | EPI_ISL_18952682 | A/Baden-Wuerttemberg/85/2024 |
| EPI_ISL_18952680 | A/Nordrhein-Westfalen/20/2024 | EPI_ISL_18941647 | A/Hessen/41/2024 |
| EPI_ISL_18952679 | A/Saarland/9/2024 | EPI_ISL_18941646 | A/Niedersachsen/65/2024 |
| EPI_ISL_18942341 | A/Nordrhein-Westfalen/18/2024 | EPI_ISL_18941645 | A/Baden-Wuerttemberg/76/2024 |
| EPI_ISL_18942340 | A/Niedersachsen/67/2024 | EPI_ISL_18941644 | A/Rheinland-Pfalz/18/2024 |
| EPI_ISL_18942339 | A/Baden-Wuerttemberg/82/2024 | EPI_ISL_18942338 | A/Rheinland-Pfalz/5/2024 |

Table 1S. Accession number (Gisaid) of influenza A(H1N1)pdm09 viruses isolated in Germany between 2009 and 2024

| | | | |
|---|---|---|---|
| EPI_ISL_18941650 | A/Nordrhein-Westfalen/46/2024 | *EPI_ISL_18941642* | *A/Baden-Wuerttemberg/75/2024* |
| EPI_ISL_18942337 | A/Nordrhein-Westfalen/47/2024 | EPI_ISL_19179543 | A/Baden-Wuerttemberg/75/2024*(cell-cultured) |
| EPI_ISL_18942336 | A/Thueringen/6/2024 | EPI_ISL_18941641 | A/Hessen/40/2024 |
| EPI_ISL_18942334 | A/Schleswig-Holstein/27/2024 | EPI_ISL_18941640 | A/Baden-Wuerttemberg/74/2024 |
| EPI_ISL_18942333 | A/Schleswig-Holstein/26/2024 | EPI_ISL_18941639 | A/Hessen/39/2024 |
| EPI_ISL_18941825 | A/Hessen/35/2024 | EPI_ISL_18941638 | A/Hessen/38/2024 |
| EPI_ISL_18941740 | A/Hessen/45/2024 | EPI_ISL_18941637 | A/Hessen/37/2024 |
| EPI_ISL_18941667 | A/Hessen/44/2024 | EPI_ISL_18941636 | A/Hessen/36/2024 |
| EPI_ISL_18941666 | A/Rheinland-Pfalz/20/2024 | EPI_ISL_18941635 | A/Bayern/35/2024 |
| EPI_ISL_18941665 | A/Rheinland-Pfalz/19/2024 | EPI_ISL_18941634 | A/Bayern/34/2024 |
| EPI_ISL_18941664 | A/Brandenburg/21/2024 | EPI_ISL_18941633 | A/Rheinland-Pfalz/17/2024 |
| EPI_ISL_18941663 | A/Brandenburg/20/2024 | EPI_ISL_18941632 | A/Sachsen/8/2024 |
| EPI_ISL_18941662 | A/Baden-Wuerttemberg/81/2024 | EPI_ISL_18941631 | A/Thueringen/39/2024 |
| EPI_ISL_18941661 | A/Baden-Wuerttemberg/80/2024 | EPI_ISL_18941630 | A/Nordrhein-Westfalen/45/2024 |
| EPI_ISL_18941660 | A/Baden-Wuerttemberg/79/2024 | EPI_ISL_18941629 | A/Thueringen/38/2024 |
| EPI_ISL_18941659 | A/Hessen/43/2024 | EPI_ISL_18941628 | A/Niedersachsen/64/2024 |
| EPI_ISL_18941658 | A/Bayern/38/2024 | EPI_ISL_18941627 | A/Niedersachsen/63/2024 |
| EPI_ISL_18941657 | A/Sachsen/18/2024 | EPI_ISL_18941626 | A/Bayern/33/2024 |
| EPI_ISL_18941656 | A/Sachsen/17/2024 | EPI_ISL_18941625 | A/Thueringen/37/2024 |
| EPI_ISL_18941655 | A/Bayern/37/2024 | EPI_ISL_18941624 | A/Bayern/32/2024 |
| EPI_ISL_18941654 | A/Bayern/36/2024 | EPI_ISL_18941623 | A/Brandenburg/19/2024 |
| EPI_ISL_18941653 | A/Niedersachsen/66/2024 | EPI_ISL_18941622 | A/Brandenburg/18/2024 |
| EPI_ISL_18941652 | A/Baden-Wuerttemberg/78/2024 | EPI_ISL_18941621 | A/Thueringen/36/2024 |
| EPI_ISL_18941651 | A/Baden-Wuerttemberg/77/2024 | EPI_ISL_18926982 | A/Nordrhein-Westfalen/36/2024 |
| EPI_ISL_18900973 | A/Baden-Wuerttemberg/51/2024 | EPI_ISL_18926981 | A/Niedersachsen/62/2024 |
| EPI_ISL_18941649 | A/Hessen/13/2024 | EPI_ISL_18926980 | A/Thueringen/26/2024 |
| EPI_ISL_18941648 | A/Hessen/42/2024 | EPI_ISL_18926979 | A/Nordrhein-Westfalen/29/2024 |
| EPI_ISL_18926977 | A/Schleswig-Holstein/21/2024 | EPI_ISL_18926978 | A/Schleswig-Holstein/22/2024 |
| EPI_ISL_18926976 | A/Schleswig-Holstein/20/2024 | EPI_ISL_18900976 | A/Thueringen/11/2024 |
| EPI_ISL_18926975 | A/Schleswig-Holstein/19/2024 | EPI_ISL_18900975 | A/Hessen/24/2024 |
| EPI_ISL_18926974 | A/Schleswig-Holstein/18/2024 | EPI_ISL_18900974 | A/Baden-Wuerttemberg/8/2024 |
| EPI_ISL_18926973 | A/Schleswig-Holstein/17/2024 | EPI_ISL_18900972 | A/Baden-Wuerttemberg/50/2024 |

Table 1S. Accession number (Gisaid) of influenza A(H1N1)pdm09 viruses isolated in Germany between 2009 and 2024

| | | | |
|---|---|---|---|
| EPI_ISL_18926972 | A/Nordrhein-Westfalen/35/2024 | EPI_ISL_18887039 | A/Rheinland-Pfalz/9/2024 |
| EPI_ISL_18926971 | A/Rheinland-Pfalz/14/2024 | EPI_ISL_18900971 | A/Sachsen-Anhalt/16/2024 |
| EPI_ISL_18926970 | A/Nordrhein-Westfalen/34/2024 | EPI_ISL_18900970 | A/Niedersachsen/38/2024 |
| EPI_ISL_18926969 | A/Nordrhein-Westfalen/33/2024 | EPI_ISL_18900969 | A/Niedersachsen/37/2024 |
| EPI_ISL_18926968 | A/Hessen/10/2024 | EPI_ISL_18900968 | A/Brandenburg/16/2024 |
| EPI_ISL_18926967 | A/Bremen/8/2024 | EPI_ISL_18900967 | A/Baden-Wuerttemberg/49/2024 |
| EPI_ISL_18926966 | A/Bremen/6/2024 | EPI_ISL_18900966 | A/Brandenburg/15/2024 |
| EPI_ISL_18926965 | A/Baden-Wuerttemberg/64/2024 | EPI_ISL_18900965 | A/Brandenburg/14/2024 |
| EPI_ISL_18926964 | A/Niedersachsen/61/2024 | EPI_ISL_18900964 | A/Schleswig-Holstein/2/2024 |
| EPI_ISL_18926963 | A/Bayern/9/2024 | EPI_ISL_18900963 | A/Nordrhein-Westfalen/31/2024 |
| EPI_ISL_18926962 | A/Niedersachsen/60/2024 | EPI_ISL_18900962 | A/Hessen/23/2024 |
| EPI_ISL_18926961 | A/Niedersachsen/59/2024 | EPI_ISL_18900961 | A/Bayern/25/2024 |
| EPI_ISL_18926960 | A/Bayern/30/2024 | EPI_ISL_18900960 | A/Sachsen/6/2024 |
| EPI_ISL_18926959 | A/Baden-Wuerttemberg/10/2024 | EPI_ISL_18900959 | A/Saarland/6/2024 |
| EPI_ISL_18926958 | A/Bayern/29/2024 | EPI_ISL_18900958 | A/Baden-Wuerttemberg/48/2024 |
| EPI_ISL_18926957 | A/Hessen/30/2024 | EPI_ISL_18900957 | A/Mecklenburg-Vorpommern/4/2024 |
| EPI_ISL_18926956 | A/Bayern/28/2024 | EPI_ISL_18900956 | A/Berlin/9/2024 |
| EPI_ISL_18926955 | A/Hessen/29/2024 | EPI_ISL_18900955 | A/Baden-Wuerttemberg/47/2024 |
| EPI_ISL_18926954 | A/Hessen/28/2024 | EPI_ISL_18900954 | A/Bremen/3/2024 |
| EPI_ISL_18926953 | A/Bayern/27/2024 | EPI_ISL_18900953 | A/Baden-Wuerttemberg/46/2024 |
| EPI_ISL_18926952 | A/Niedersachsen/20/2024 | EPI_ISL_18900952 | A/Hessen/7/2024 |
| EPI_ISL_18926951 | A/Nordrhein-Westfalen/19/2024 | EPI_ISL_18900951 | A/Bayern/24/2024 |
| EPI_ISL_18900981 | A/Berlin/13/2024 | EPI_ISL_18900950 | A/Brandenburg/7/2024 |
| *EPI_ISL_18514885* | *A/Niedersachsen/22/2023* | EPI_ISL_18900949 | A/Baden-Wuerttemberg/45/2024 |
| EPI_ISL_19179546 | A/Niedersachsen/22/2023*(cell-cultured) | EPI_ISL_18900948 | A/Thueringen/4/2024 |
| EPI_ISL_18900979 | A/Rheinland-Pfalz/13/2024 | EPI_ISL_18900947 | A/Nordrhein-Westfalen/30/2024 |
| EPI_ISL_18900978 | A/Hessen/25/2024 | EPI_ISL_18900946 | A/Bayern/23/2024 |
| EPI_ISL_18900977 | A/Bremen/13/2024 | EPI_ISL_18900945 | A/Bayern/22/2024 |
| EPI_ISL_18900944 | A/Niedersachsen/9/2024 | EPI_ISL_18887042 | A/Schleswig-Holstein/8/2024 |
| EPI_ISL_18887073 | A/Baden-Wuerttemberg/43/2024 | EPI_ISL_18887041 | A/Rheinland-Pfalz/10/2024 |
| EPI_ISL_18887072 | A/Baden-Wuerttemberg/42/2024 | EPI_ISL_18887040 | A/Rheinland-Pfalz/3/2024 |
| EPI_ISL_18887071 | A/Hessen/21/2024 | EPI_ISL_18887038 | A/Bayern/18/2024 |

Table 1S. Accession number (Gisaid) of influenza A(H1N1)pdm09 viruses isolated in Germany between 2009 and 2024

| | | | |
|---|---|---|---|
| EPI_ISL_18887070 | A/Bayern/19/2024 | EPI_ISL_18869445 | A/Brandenburg/1/2024 |
| EPI_ISL_18887069 | A/Baden-Wuerttemberg/41/2024 | EPI_ISL_18887037 | A/Bayern/17/2024 |
| EPI_ISL_18887068 | A/Rheinland-Pfalz/11/2024 | EPI_ISL_18885546 | A/Nordrhein-Westfalen/22/2024 |
| EPI_ISL_18887067 | A/Baden-Wuerttemberg/7/2024 | EPI_ISL_18882014 | A/Bayern/3/2024 |
| EPI_ISL_18887066 | A/Nordrhein-Westfalen/27/2024 | EPI_ISL_18882013 | A/Bayern/16/2024 |
| EPI_ISL_18887065 | A/Baden-Wuerttemberg/40/2024 | EPI_ISL_18882012 | A/Bayern/15/2024 |
| EPI_ISL_18887064 | A/Hessen/20/2024 | EPI_ISL_18882011 | A/Bayern/14/2024 |
| EPI_ISL_18887063 | A/Baden-Wuerttemberg/39/2024 | EPI_ISL_18882010 | A/Rheinland-Pfalz/2/2024 |
| EPI_ISL_18887062 | A/Hessen/19/2024 | EPI_ISL_18882009 | A/Hessen/15/2024 |
| EPI_ISL_18887061 | A/Nordrhein-Westfalen/26/2024 | EPI_ISL_18882008 | A/Bayern/2/2024 |
| EPI_ISL_18887060 | A/Hessen/18/2024 | EPI_ISL_18882007 | A/Bayern/13/2024 |
| EPI_ISL_18887059 | A/Nordrhein-Westfalen/8/2024 | EPI_ISL_18882006 | A/Bremen/11/2024 |
| EPI_ISL_18887058 | A/Nordrhein-Westfalen/25/2024 | EPI_ISL_18882005 | A/Bremen/9/2024 |
| EPI_ISL_18887057 | A/Nordrhein-Westfalen/24/2024 | EPI_ISL_18882004 | A/Baden-Wuerttemberg/27/2024 |
| EPI_ISL_18887056 | A/Nordrhein-Westfalen/23/2024 | EPI_ISL_18882003 | A/Brandenburg/12/2024 |
| EPI_ISL_18887055 | A/Brandenburg/6/2024 | EPI_ISL_18882002 | A/Hessen/14/2024 |
| EPI_ISL_18887054 | A/Hessen/17/2024 | EPI_ISL_18882001 | A/Bayern/12/2024 |
| EPI_ISL_18887053 | A/Berlin/20/2024 | EPI_ISL_18882000 | A/Thueringen/9/2024 |
| EPI_ISL_18887052 | A/Berlin/7/2024 | EPI_ISL_18881999 | A/Baden-Wuerttemberg/3/2024 |
| EPI_ISL_18887051 | A/Schleswig-Holstein/10/2023 | EPI_ISL_18881998 | A/Brandenburg/11/2024 |
| EPI_ISL_18887050 | A/Schleswig-Holstein/15/2024 | EPI_ISL_18881997 | A/Nordrhein-Westfalen/7/2024 |
| EPI_ISL_18887049 | A/Schleswig-Holstein/14/2024 | EPI_ISL_18881996 | A/Nordrhein-Westfalen/5/2024 |
| EPI_ISL_18887048 | A/Schleswig-Holstein/13/2024 | EPI_ISL_18881995 | A/Baden-Wuerttemberg/26/2024 |
| EPI_ISL_18887047 | A/Schleswig-Holstein/9/2023 | EPI_ISL_18881994 | A/Brandenburg/14/2023 |
| EPI_ISL_18887046 | A/Schleswig-Holstein/12/2024 | EPI_ISL_18881993 | A/Brandenburg/13/2023 |
| EPI_ISL_18887045 | A/Schleswig-Holstein/11/2024 | EPI_ISL_18881992 | A/Nordrhein-Westfalen/3/2024 |
| EPI_ISL_18887044 | A/Schleswig-Holstein/10/2024 | EPI_ISL_18881991 | A/Saarland/1/2024 |
| EPI_ISL_18887043 | A/Schleswig-Holstein/9/2024 | EPI_ISL_18881990 | A/Sachsen/2/2024 |
| EPI_ISL_18881989 | A/Baden-Wuerttemberg/50/2023 | EPI_ISL_18869448 | A/Nordrhein-Westfalen/15/2024 |
| EPI_ISL_18881988 | A/Baden-Wuerttemberg/49/2023 | EPI_ISL_18869447 | A/Rheinland-Pfalz/1/2024 |
| EPI_ISL_18881987 | A/Baden-Wuerttemberg/48/2023 | EPI_ISL_18869446 | A/Berlin/2/2024 |
| EPI_ISL_18881986 | A/Brandenburg/12/2023 | EPI_ISL_18869444 | A/Sachsen-Anhalt/1/2024 |

Table 1S. Accession number (Gisaid) of influenza A(H1N1)pdm09 viruses isolated in Germany between 2009 and 2024

| | | | |
|---|---|---|---|
| EPI_ISL_18881985 | A/Berlin/46/2023 | EPI_ISL_18831405 | A/Berlin/58/2023 |
| EPI_ISL_18881984 | A/Baden-Wuerttemberg/47/2023 | EPI_ISL_18869443 | A/Baden-Wuerttemberg/21/2024 |
| EPI_ISL_18881983 | A/Brandenburg/11/2023 | EPI_ISL_18869442 | A/Baden-Wuerttemberg/1/2024 |
| EPI_ISL_18881982 | A/Rheinland-Pfalz/14/2023 | EPI_ISL_18869441 | A/Berlin/66/2023 |
| EPI_ISL_18881981 | A/Nordrhein-Westfalen/24/2023 | EPI_ISL_18869440 | A/Berlin/48/2023 |
| EPI_ISL_18881980 | A/Baden-Wuerttemberg/46/2023 | EPI_ISL_18869439 | A/Niedersachsen/39/2023 |
| EPI_ISL_18881979 | A/Berlin/67/2023 | EPI_ISL_18839331 | A/Mecklenburg-Vorpommern/16/2023 |
| EPI_ISL_18869469 | A/Niedersachsen/14/2024 | EPI_ISL_18839330 | A/Mecklenburg-Vorpommern/15/2023 |
| EPI_ISL_18869468 | A/Baden-Wuerttemberg/25/2024 | EPI_ISL_18839329 | A/Mecklenburg-Vorpommern/14/2023 |
| EPI_ISL_18869467 | A/Saarland/3/2024 | EPI_ISL_18839328 | A/Bayern/20/2023 |
| EPI_ISL_18869466 | A/Brandenburg/10/2024 | EPI_ISL_18839327 | A/Sachsen/5/2023 |
| EPI_ISL_18869465 | A/Brandenburg/9/2024 | EPI_ISL_18839326 | A/Sachsen/6/2023 |
| EPI_ISL_18869464 | A/Sachsen/3/2024 | EPI_ISL_18839325 | A/Hessen/44/2023 |
| EPI_ISL_18869463 | A/Nordrhein-Westfalen/17/2024 | EPI_ISL_18839324 | A/Hessen/43/2023 |
| EPI_ISL_18869462 | A/Hessen/12/2024 | EPI_ISL_18839323 | A/Baden-Wuerttemberg/45/2023 |
| EPI_ISL_18869461 | A/Baden-Wuerttemberg/24/2024 | EPI_ISL_18839322 | A/Baden-Wuerttemberg/44/2023 |
| EPI_ISL_18869460 | A/Baden-Wuerttemberg/23/2024 | EPI_ISL_18839321 | A/Saarland/5/2023 |
| EPI_ISL_18869459 | A/Thueringen/1/2024 | EPI_ISL_18839320 | A/Bayern/14/2023 |
| EPI_ISL_18869458 | A/Bremen/7/2024 | EPI_ISL_18839319 | A/Hessen/42/2023 |
| EPI_ISL_18869457 | A/Brandenburg/2/2024 | EPI_ISL_18839318 | A/Mecklenburg-Vorpommern/10/2023 |
| EPI_ISL_18869456 | A/Sachsen-Anhalt/2/2024 | EPI_ISL_18839317 | A/Mecklenburg-Vorpommern/13/2023 |
| EPI_ISL_18869455 | A/Bayern/11/2024 | EPI_ISL_18839316 | A/Mecklenburg-Vorpommern/12/2023 |
| EPI_ISL_18869454 | A/Hessen/11/2024 | EPI_ISL_18839315 | A/Mecklenburg-Vorpommern/9/2023 |
| EPI_ISL_18869453 | A/Bayern/10/2024 | EPI_ISL_18839314 | A/Bremen/4/2023 |
| EPI_ISL_18869452 | A/Nordrhein-Westfalen/16/2024 | EPI_ISL_18839313 | A/Thueringen/12/2023 |
| EPI_ISL_18869451 | A/Baden-Wuerttemberg/38/2023 | EPI_ISL_18839312 | A/Baden-Wuerttemberg/43/2023 |
| EPI_ISL_18869450 | A/Baden-Wuerttemberg/22/2024 | EPI_ISL_18839311 | A/Berlin/45/2023 |
| EPI_ISL_18869449 | A/Hessen/1/2024 | EPI_ISL_18839310 | A/Sachsen-Anhalt/14/2023 |
| EPI_ISL_18839309 | A/Sachsen-Anhalt/12/2023 | EPI_ISL_18831408 | A/Berlin/60/2023 |
| EPI_ISL_18839308 | A/Hessen/41/2023 | EPI_ISL_18831407 | A/Berlin/59/2023 |
| EPI_ISL_18839307 | A/Hessen/33/2023 | EPI_ISL_18831406 | A/Hessen/31/2023 |
| EPI_ISL_18839306 | A/Sachsen/4/2023 | EPI_ISL_18831404 | A/Berlin/57/2023 |

Table 1S. Accession number (Gisaid) of influenza A(H1N1)pdm09 viruses isolated in Germany between 2009 and 2024

| | | | |
|---|---|---|---|
| EPI_ISL_18839305 | A/Baden-Wuerttemberg/42/2023 | EPI_ISL_18576678 | A/Mecklenburg-Vorpommern/4/2023 |
| EPI_ISL_18839304 | A/Baden-Wuerttemberg/33/2023 | EPI_ISL_18831403 | A/Berlin/56/2023 |
| EPI_ISL_18839303 | A/Rheinland-Pfalz/12/2023 | EPI_ISL_18831402 | A/Niedersachsen/24/2023 |
| EPI_ISL_18839302 | A/Baden-Wuerttemberg/41/2023 | EPI_ISL_18831401 | A/Berlin/42/2023 |
| EPI_ISL_18839301 | A/Brandenburg/9/2023 | EPI_ISL_18831400 | A/Brandenburg/8/2023 |
| EPI_ISL_18839300 | A/Rheinland-Pfalz/15/2023 | EPI_ISL_18831399 | A/Baden-Wuerttemberg/31/2023 |
| EPI_ISL_18839299 | A/Sachsen-Anhalt/13/2023 | EPI_ISL_18831398 | A/Bayern/19/2023 |
| EPI_ISL_18839298 | A/Berlin/65/2023 | EPI_ISL_18831397 | A/Rheinland-Pfalz/10/2023 |
| EPI_ISL_18839297 | A/Berlin/64/2023 | EPI_ISL_18831396 | A/Brandenburg/7/2023 |
| EPI_ISL_18839296 | A/Berlin/63/2023 | EPI_ISL_18831395 | A/Hessen/29/2023 |
| EPI_ISL_18839295 | A/Nordrhein-Westfalen/20/2023 | EPI_ISL_18831394 | A/Hessen/28/2023 |
| EPI_ISL_18839294 | A/Bremen/5/2023 | EPI_ISL_18831393 | A/Hessen/27/2023 |
| EPI_ISL_18839293 | A/Berlin/62/2023 | EPI_ISL_18831392 | A/Hessen/26/2023 |
| EPI_ISL_18839292 | A/Baden-Wuerttemberg/40/2023 | *EPI_ISL_18831390* | *A/Saarland/3/2023* |
| EPI_ISL_18839291 | A/Hessen/40/2023 | EPI_ISL_19179545 | A/Saarland/3/2023* (cell-cultured) |
| EPI_ISL_18832503 | A/Berlin/50/2023 | EPI_ISL_18831389 | A/Berlin/41/2023 |
| EPI_ISL_18831420 | A/Berlin/51/2023 | EPI_ISL_18831388 | A/Niedersachsen/37/2023 |
| EPI_ISL_18831419 | A/Berlin/52/2023 | EPI_ISL_18788624 | A/Hessen/25/2023 |
| EPI_ISL_18831418 | A/Berlin/53/2023 | EPI_ISL_18788621 | A/Brandenburg/6/2023 |
| EPI_ISL_18831417 | A/Hessen/35/2023 | EPI_ISL_18788619 | A/Brandenburg/5/2023 |
| EPI_ISL_18831416 | A/Hessen/36/2023 | EPI_ISL_18788617 | A/Hessen/23/2023 |
| EPI_ISL_18831415 | A/Hessen/37/2023 | EPI_ISL_18788616 | A/Thueringen/9/2023 |
| EPI_ISL_18831414 | A/Hessen/38/2023 | EPI_ISL_18788615 | A/Niedersachsen/25/2023 |
| EPI_ISL_18831413 | A/Bremen/3/2023 | EPI_ISL_18788614 | A/Berlin/49/2023 |
| EPI_ISL_18831412 | A/Bremen/2/2023 | EPI_ISL_18788613 | A/Thueringen/8/2023 |
| EPI_ISL_18831411 | A/Bremen/6/2023 | EPI_ISL_18788612 | A/Baden-Wuerttemberg/39/2023 |
| EPI_ISL_18831410 | A/Nordrhein-Westfalen/15/2023 | EPI_ISL_18788611 | A/Rheinland-Pfalz/7/2023 |
| EPI_ISL_18831409 | A/Hessen/39/2023 | EPI_ISL_18651955 | A/Berlin/29/2023 |
| EPI_ISL_18788609 | A/Niedersachsen/23/2023 | EPI_ISL_18651954 | A/Berlin/30/2023 |
| EPI_ISL_18651965 | A/Berlin/39/2023 | EPI_ISL_18651953 | A/Brandenburg/4/2023 |
| EPI_ISL_18651964 | A/Berlin/34/2023 | EPI_ISL_18576679 | A/Nordrhein-Westfalen/11/2023 |
| EPI_ISL_18651963 | A/Berlin/36/2023 | EPI_ISL_18576677 | A/Schleswig-Holstein/5/2023 |

Table 1S. Accession number (Gisaid) of influenza A(H1N1)pdm09 viruses isolated in Germany between 2009 and 2024

| | | | |
|---|---|---|---|
| EPI_ISL_18651962 | A/Berlin/35/2023 | EPI_ISL_18651956 | A/Berlin/31/2023 |
| EPI_ISL_18651961 | A/Mecklenburg-Vorpommern/5/2023 | *EPI_ISL_18900980* | *A/Niedersachsen/39/2024* |
| EPI_ISL_18651960 | A/Rheinland-Pfalz/8/2023 | EPI_ISL_19179544 | A/Niedersachsen/39/2024* (cell-cultured) |
| EPI_ISL_18651959 | A/Berlin/33/2023 | EPI_ISL_18514884 | A/Berlin/26/2023 |
| EPI_ISL_18651957 | A/Berlin/38/2023 | EPI_ISL_18788610 | A/Saarland/4/2023 |
